# Unveiling G-Protein-Coupled Receptor Conformational Dynamics via Metadynamics Simulations and Markov State Models

**DOI:** 10.1101/2025.01.21.634062

**Authors:** Rita A. Roessner, Nicolas Floquet, Maxime Louet

## Abstract

The dynamic character of G-protein-coupled receptors (GPCRs) is essential in their functionality as signal transducers. However, the molecular details of how ligands affect this conformational repertoire to steer intracellular signaling pathways remain elusive. Here, we address this question by modeling the conformational landscape of the growth hormone secretagogue receptor (GHSR-1a), a prototypical peptide-activated class A GPCR, in its apo state and bound to pharmacologically distinct ligands. We present a generally applicable protocol to efficiently explore the conformational space of GHSR-1a that is sensitive to the bound ligand. Combining metadynamics simulations and Markov state modeling, we computed the free energy landscape of GHSR-1a in its apo state and bound to an agonist, antagonist, or inverse agonist, respectively. Consistent with the current multi-state model of GPCR activity, we found that GHSR-1a populates multiple metastable states whose energies and transition probabilities change depending on the bound ligand. Furthermore, our Markov state models (MSM) have revealed two intermediate states that have not yet been described by experimental structures and which we assume to facilitate the binding of extracellular ligands and intracellular protein partners, respectively. Lastly, our MSMs allowed us to shed light on the molecular differences between basal and agonist-induced GHSR-1a activation. Our results are not only compatible with previously reported experimental data, but they capture the equilibria governing GHSR-1a activation in unprecedented detail. Due to its applicability to all class A GPCRs, our protocol is a valuable tool for the development of pharmaceuticals targeting this protein family.

## INTRODUCTION

G-protein-coupled receptors (GPCRs) are the most important family of membrane proteins responsible for cellular responses to a variety of different stimuli, including hormones, neurotransmitters, and exogenous substances. It is estimated that one-third of approved pharmaceutical drugs target GPCRs [1]. Understanding the dynamics of GPCR-mediated signaling at high resolution has consequently been a longstanding goal toward designing more effective and less toxic drugs. All rhodopsin-like, class-A GPCRs exhibit a common structure comprising seven transmembrane (TM) helices (Fig. 1A), which define a binding pocket able to recognize extracellular ligands. The binding of ligands into this well-defined pocket is the first step to promoting coupling and activation of intracellular protein partners (transducers), including G-proteins, GPCRs kinases, and arrestins [2]. Over the past decade, the number of structures describing these receptors has grown exponentially, exceeding 1000 structures to date [3]. Even though these experimental structures have provided invaluable insight into the conformational plasticity of GPCRs, they only capture a few distinct lowest-energy states that do not allow a comprehensive understanding of the allosteric control of GPCR activation by extracellular ligands. Indeed, biophysical studies point to a mechanism where GPCR signaling is governed by a spectrum of conformational states in constant dynamic equilibrium. In this context, extracellular ligands and intracellular protein partners modulate these equilibria, resulting in complex energy landscapes [4], [5].

**Figure 1:**
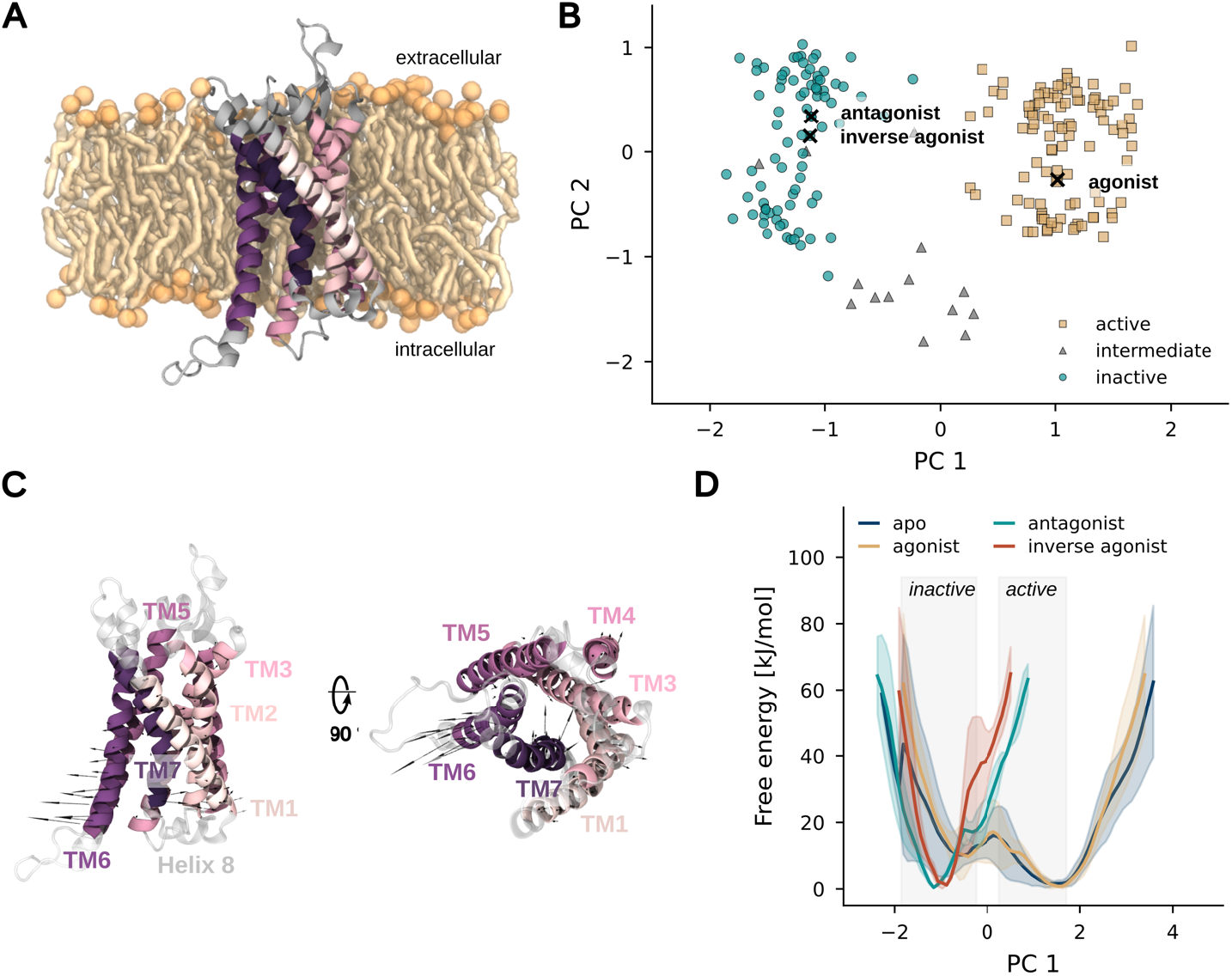
PCA-based metadynamics simulations of GHSR-1a. (A) Side-cut view of GHSR-1a (cartoon representation) embedded in a POPC membrane. The residues of the membrane-spanning region of the receptor considered in PCA are colored by index (pink to purple from the N-to the C-terminus); the remaining residues are shown in gray. Lipids are depicted in orange, their head group phosphorus atoms being represented as spheres whereas lipid tails were reported as sticks. (B) Projection of the experimental class A GPCR structures considered for PCA onto the first two PCs. Each marker represents a structure that was annotated as active (yellow squares), inactive (green circles), or intermediate (gray triangles) in GPCRdb. Relevant GHSR-1a structures are highlighted with black crosses (together with the bound ligand type). (C) Side and top view of GHSR-1a (colored as in A). The arrows indicate the collective motion along PC1, with the arrow length corresponding to the magnitude of the vector. (D) Free energy profiles (average ± standard deviation, n=3 independent simulation sets) as a function of PC1 for the apo (blue, PDB id. 7F9Y), agonist-bound (yellow, PDB id. 7F9Y), antagonist-bound (green, PDB id. 6KO5), and inverse agonist-bound red, PDB id. 7F83) systems. The gray boxes represent the range of the projection values of active and inactive experimental structures.

Classical Molecular dynamics (MD) simulations could appear as a method of choice to bridge the gap between structures and biophysical studies [6]. However, although technical advances in computer architecture have considerably increased the lengths of state-of-the-art trajectories, observing and discussing the activation/inhibition of these membrane receptors at the molecular scale still requires the use of enhanced sampling protocols. This is particularly true when one wants to compute a free energy landscape for these receptors, a task requiring capturing a significant number of transitions between all concerned conformational states. Thus, approaches based on collective variables (CV), such as metadynamics have been extensively employed over recent years to shed light on the energy landscape of GPCRs and its modulation by ligands [7], [8], [9], [10]. In such studies, the main challenge most often resides in identifying CVs that can clearly distinguish the relevant conformational states and enhance the sampling of the slowest degrees of freedom involved in the process of interest [11], [12]. Among other studies, CVs leveraged in recent works range from the orientation of certain residues known to be relevant in the GPCR activation process, so-called microswitches [8], the RMSD to a reference structure (most often active or inactive ones) [9], or a set of interhelical distances linearly combined into a “general activation index” [10]. Whereas these variables are indeed capable of giving rise to a convergenced free energy profile, it is often at the expense of their ability to discriminate accurately between the explored conformations (several, completely different conformations can display the same CV value). Additionally, they often neglect collective motions which are crucial to sample the large conformational transitions GPCRs undergo.

In this study, we implemented a computational strategy combining biased (metadynamics) and unbiased molecular dynamics simulations as well as Markov state modeling to characterize the conformational landscape of the growth hormone secretagogue receptor (GHSR-1a). Even though GHSR-1a may be described as a prototypical class A GPCR it exhibits substantial constitutive activity (= activity without a ligand bound), qualifying it as an excellent model system to study the activation and inhibition of GPCRs. Additionally, several structures of the receptor bound to various ligands, which include members of different classes such as agonists [13], [14], antagonists [15], and inverse agonists [16], have been recently resolved. Here, we demonstrate that a principal component analysis (PCA) of experimental class A GPCR structures is an efficient way to derive good CVs for metadynamics simulations generating representative sets of initial conformations required for the efficient construction of Markov state models. Notably, by using our general CVs as input to simulations of other class A GPCRs, our approach can likely be transferred to other receptors of that protein family. Furthermore, our data resulting from Markovian analyses of an extensive set of free MD simulations show a clear ligand-induced kinetic and thermodynamic remodeling of the equilibria that constitute the conformational landscape of GHSR-1a.

## METHODS

The principal component analysis (PCA) was performed with GROMACS tools [18]. The PDB files of 612 published class A GPCR structures (state: 11/2022, S1 Table) were processed to obtain solely the Cα coordinates of the receptor chain [3]. To acquire a set of equivalent Cα atoms on which to perform PCA, we performed sequence-based alignment of all structures utilizing the GPCRdb numbering scheme [19]. Among these 612 structures, 107 lacked important residues in the 7TM core, thus we chose to discard them to maintain a good balance between the structural variability (i.e. maximizing the number of structures) and the structural description (i.e. maximizing the number of Cα considered). 505 structures were kept with the seven-helical core region largely conserved (186 Cα atoms - Fig. 1A). We could not consider loop regions due to their high sequence variability among class A GPCRs that did not allow a good sequence alignment. Additionally, because of their high flexibility loops are often not resolved in experimental structures. An RMSD-based clustering with a cutoff of 0.6 Å was conducted on the respective Cα atoms of this subset of structures using *gmx cluster*. The objective of this step was to remove potential biases rooted in scientific interest or preferential experimental conditions overrepresenting specific receptors or certain conformations. The conserved Cα atoms of the resulting 200 structures were subjected to PCA where the covariance matrix was calculated and diagonalized with *gmx covar* and the obtained eigenvectors were analyzed using *gmx anaeig*.

Well-tempered metadynamics simulations [20] were carried out using the PLUMED 2.8.1 plug-in [21], [22] incorporated into GROMACS [18]. As collective variables, we used the amplitudes of displacement with respect to the initial structure along the eight first eigenvectors (those with the highest variance) derived from PCA (PCAVAR descriptor implemented in PLUMED). More precisely, in every experiment, eight replicas were run in parallel with a bias-exchange scheme [23]. Each simulation was biased along a different single CV, being the amplitude of displacement along a particular eigenvector, and exchanged regularly with other replicas. For each system, we conducted n=3 independent experiments. Gaussian hills with an initial height of 1.0 kJ·mol^−1^ were deposited every 10 ps with a bias factor of 10. The Gaussian width was set to 10^−4^ based on the projection values of the experimental structures to ensure a good resolution of the resulting free energy profiles. An exchange between randomly chosen replicas was attempted every 100 ps with a success rate of ∼20 % based on the Boltzmann criterion. Free energies were calculated using the sum_hills function of the PLUMED plugin. Convergence was assessed by monitoring the time evolution of the estimated free energy as a function of the projection value along the first eigenvector (S4 Fig.). Depending on the system, the energy profiles converged after ∼8 μs (8 × 1µs) of accumulative simulation time.

To effectively sample the GHSR-1a conformational landscape with standard (unbiased) MD simulations, a finite number of diverse models was chosen from the configurations explored by the respective system in metadynamics simulations. To do so, the models obtained from metadynamics simulations were projected onto the eight eigenvectors used as collective variables. Regular space clustering performed on this data allowed the extraction of 50 representative conformations that were used to initiate the first round of 700 ns-long unbiased MD simulations with n=4 simulation repeats. Subsequently, several rounds of adaptive sampling were conducted by constructing an MSM utilizing existing unbiased MD simulations and randomly selecting microstates to sample from, based on the stationary distribution over microstates. Thereby the probability of choosing a microstate as a new initial configuration was inversely proportional to its stationary probability. Consequently, microstates with a low stationary probability were more likely to be selected for seeding new simulations which has been demonstrated to be efficient for exploring new states [24], [25]. After four rounds of such adaptive sampling, a combined simulation time of at least 160 μs was accumulated for each system. For the construction of the MSM, we considered only the last 500 ns of each simulation resulting in a total simulation time of at least 116 μs per system.

Overall, the construction of the MSMs was performed using the Python packages pyEMMA [26] and Deeptime [27] for feature scoring, data projection, building the MSM, and extracting the final macrostates. Molecular features for MSM construction were generated by calculating the 754 Cα-Cα distances between a subset of 42 Cα atoms evenly spanning the structurally conserved transmembrane region of GHSR-1a (S6 Fig.). Dimensionality reduction of the data was performed through projection onto the slowest degrees of freedom using time-lagged independent component analysis (TICA) [28] with a time-lag of 20 ns. In general, MSMs are constructed with the assumption of exhaustive sampling of the equilibrium distribution, however, previous studies point out that this is largely unfeasible for large protein systems [29]. Consequently, the variational approach for Markov processes (VAMP), generally used for MSM hyperparameter selection, tends to favor the exploration of under-sampled processes rather than the convergence of a few timescales of interest [30], [31], [32]. Following the adapted protocol of Bergh et al., we calculated the stationary probability of the lowest-energy state using PCCA+ [33] for various hyperparameter combinations [29]. We obtained consistent results performing k-means clustering on the two slowest independent components (ICs) selecting 100 clusters (S7 Fig.). To choose a Markovian lag time we estimated the implied timescales (ITS) from the transition probability matrices at different lag times [34], [35]. The uncertainty of the ITS is quantified based on Markov models sampled according to a Bayesian scheme. Since at a lag time of 100 ns the ITS were approximately invariant within the error this lag time was selected to build the Markov models (S8 Fig.). In addition to ITS, the MSMs were validated by the Chapman-Kolmogrov test [34] in which the ability of a transition matrix to reproduce transition probabilities at longer timescales is evaluated (S11 Fig.). For the sake of human interpretation, the 100 microstates were coarse-grained into two to three metastable macrostates using the PCCA+ algorithm [33] (Fig. 2B, Fig. 3A). Mean first passage times between pairs of states (MFPT_ij_) were computed based on the stationary distribution of the macrostates and the transition probability matrix of the MSM.

**Figure 2:**
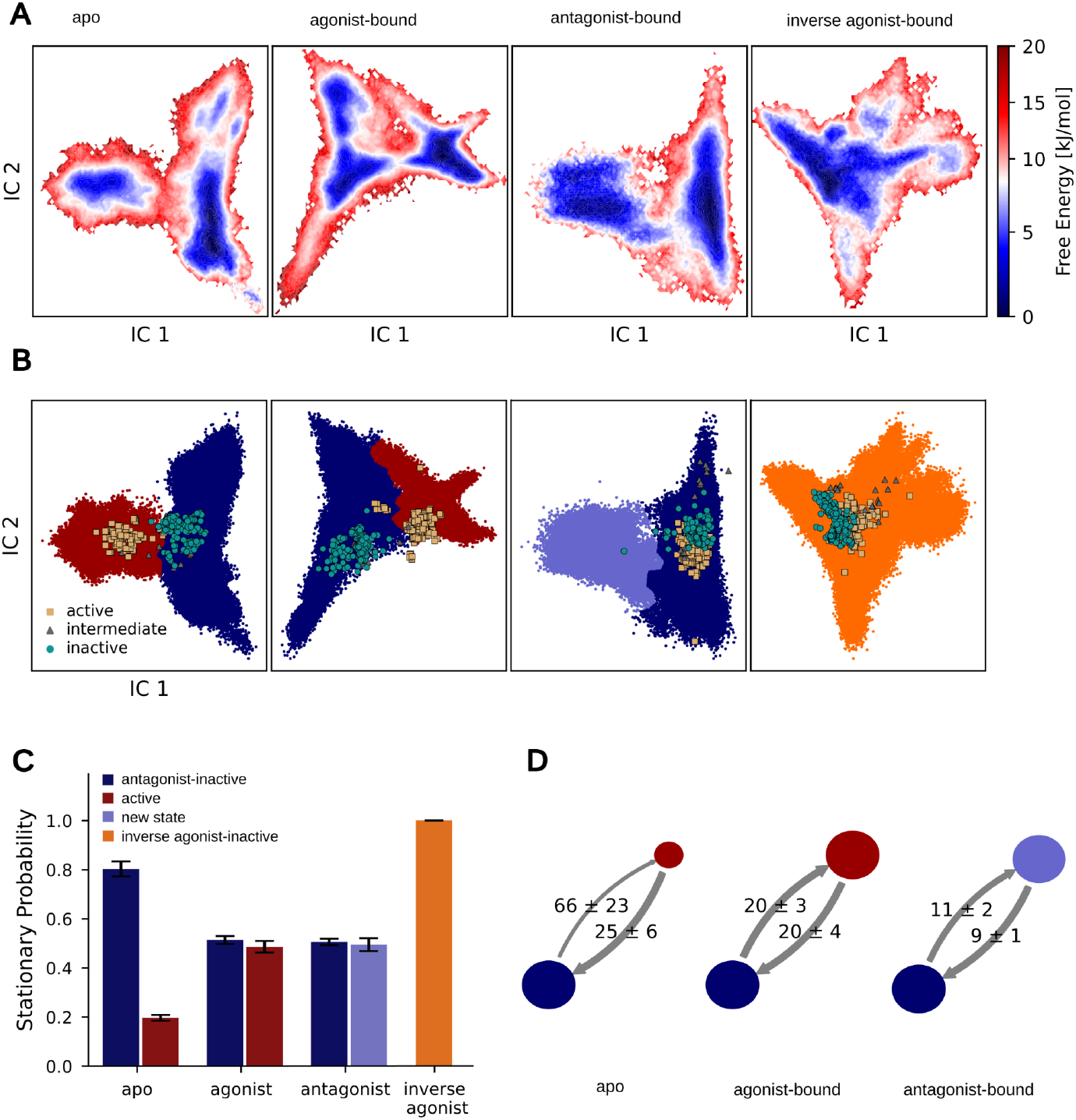
MSM free energy landscape of GHSR-1a systems. A: Projection of the MSM free energy surface onto the two slowest ICs of each system. B: For each system, the metastable states separated by the highest free energy barrier are presented in red (active), dark blue (antagonist-inactive), light blue (new state), or orange (inverse agonist-inactive). Experimental class A GPCR structures are indicated by yellow squares (active), green circles (inactive), or gray triangles (intermediate). C: Stationary probability of the two most distinct macrostates for each system. D: Mean first passage times between states are indicated with directional arrows and reported in microseconds. The disk sizes are proportional to the stationary probabilities of the active (red), inactive, 6KO5-like (blue), and the new state (light blue). All errors in (C) and (D) indicate the lower and upper bounds of the 95% confidence interval calculated from n=100 samples drawn from the Bayesian posterior distribution of the transition matrix.

**Figure 3:**
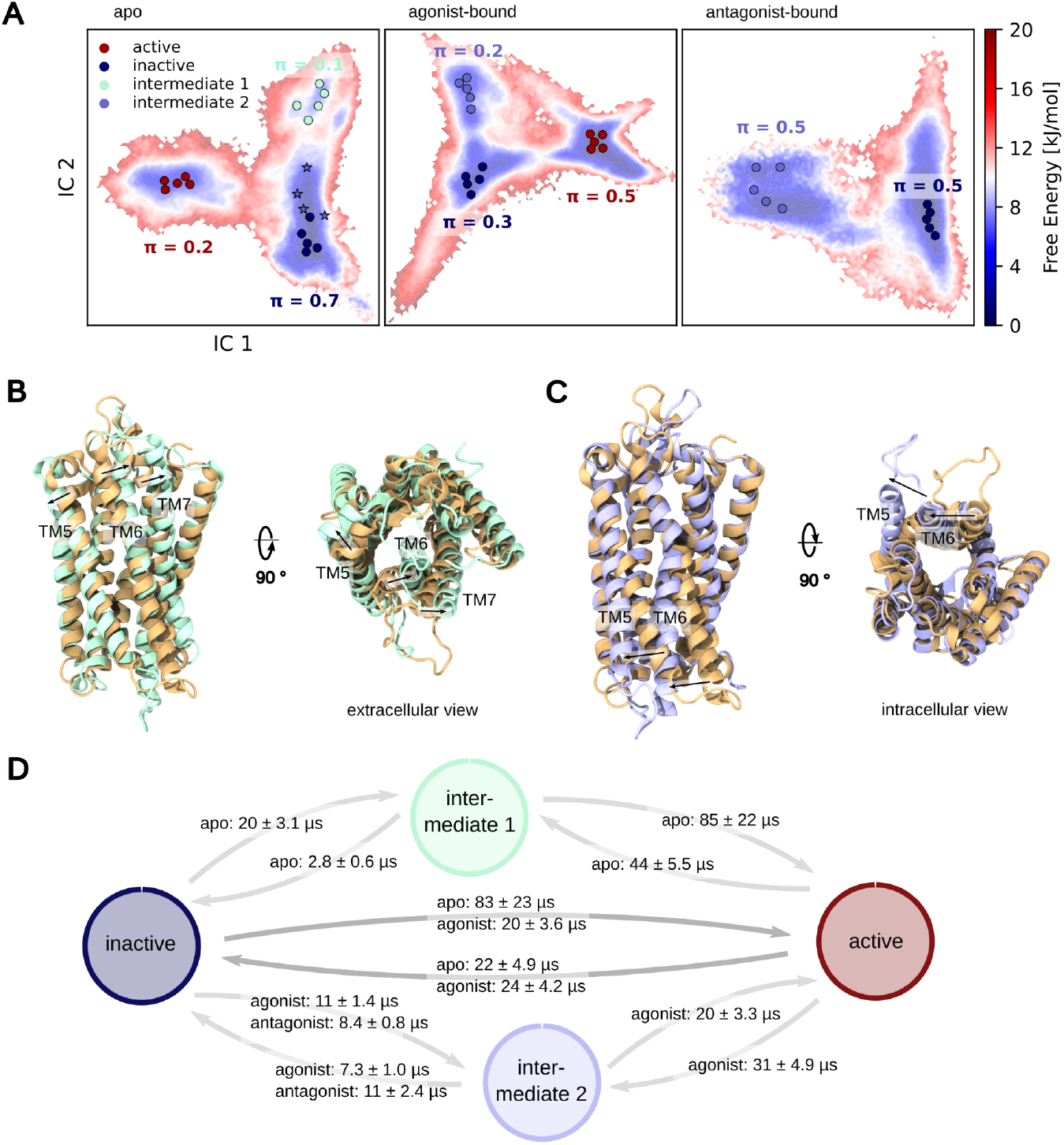
Refined clustering of GHSR-1a free energy landscapes. A: MSM free energy landscape of the apo (left), agonist-bound (middle), and antagonist-bound (right) GHSR-1a. The n=5 lowest energy models for each macrostate are highlighted in red (active), blue (inactive), light green (i*ntermediate 1*), and light blue (*intermediate 2*). The models of the apo *intermediate 2* represented as * were not assigned to the inactive macrostate during PCCA+ clustering. All macrostates are annotated with their respective stationary probability π. B: Superposition of PDB id. 7F9Y (yellow) and *intermediate 1* (light green), shown from side and extracellular views. C: Overlay of PDB id. 7F9Y (orange) and *intermediate 2* (light blue), depicted from side and intracellular views. D: Mean first passage times (MFPT) between metastable states. The errors indicate the lower and upper bounds of the 95% confidence interval calculated from n=100 samples drawn from the Bayesian posterior distribution of the transition matrix.

## RESULTS

### Eigenvectors from principal component analysis on class A GPCR structures enabled sampling of GHSR-1a activation motions

As principal component analysis (PCA) has previously been shown to be a powerful tool to unravel collective motions from experimental structures, we employed this dimensionality reduction approach to a representative subset of currently solved class A GPCR structures (S1 Table) [36]. We considered the coordinates of 186 Cα atoms largely spanning the seven-helical core region of the GPCRs (Fig. 1A) which were present in more than 80 % of the published structures [2]. To remove potential biases rooted in scientific interest or preferential experimental conditions overrepresenting certain conformations or specific receptors, we additionally performed an RMSD-based clustering on the respective Cα atoms reducing the input for PCA to a total of 200 structures (see Methods section). The projection of the experimental structures onto the first two principal components (PCs) is reported in Fig. 1B. In this 2D space, the PCA separated the structures into three clusters which largely coincided with the distance-based GPCRdb annotation for active, intermediate, and inactive conformations [19]. Furthermore, PC1 not only unambiguously distinguished between active and inactive experimental structures but also described the outward movement of TM6 with the simultaneous slight inward movement of TM7 on the intracellular side, which is a prominent feature of GPCR activation (Fig. 1C) [14], [37]. This further supported that the PCs associated with dominant eigenvectors captured collective motions relevant to the activation process of GPCRs, making them interesting collective variables (CVs) for metadynamics simulations.

To comprehensively sample the conformational space of GHSR-1a we used the first eight PCs covering 64 % of the variance encoded by the experimental data (S2 Fig., S3 Fig.) as CVs for metadynamics simulations. The main drawback of these CVs is, that they are by definition of PCA, orthogonal, and uncoupled. To overcome this limitation, we employed a bias-exchange scheme detailed in the Methods section. Consequently, one experiment entailed eight 1 µs-long replicas, each replica being restrained along one of the eight selected PCs. The eight CVs were sequentially coupled by random bias-exchange trials every 500 ps based on the Boltzmann criterion. For each system (apo, ghrelin-bound, antagonist-bound, and inverse agonist-bound GHSR-1a) we conducted n=3 independent experiments resulting in an accumulative simulation time of 24 µs per system. We confirmed the convergence of the simulations by monitoring the free energy profiles as a function of time (S4 Fig.). Based on the free energy profiles we deduced that the apo and agonist-bound systems were able to explore a broader conformational space than that described by experimental data with the minima of the double-well profile projected onto active and inactive experimental structures, respectively.

Interestingly, no transition from the inactive to the active conformation was captured for the antagonist-bound and inverse agonist-bound systems. Both systems showed a profile with a single minimum, indicating a restriction of the conformational space to inactive receptor conformations, which was more pronounced for the inverse agonist (Fig. 1D). As the same protocol was applied to all systems, our results suggest that the energy barrier separating active and inactive conformations could be harder to cross in the presence of the antagonist or the inverse agonist compared to the receptor in its apo or agonist-bound state. In agreement with previous experimental observations, this conformational restriction could be interpreted as a stabilization effect of the inactive conformation by the antagonist and inverse agonist molecules in the absence of the G-protein [15], [16].

Taken together this indicates that our protocol was successful in capturing the effect of pharmacologically distinct ligands. However, it failed to quantitatively assess the height of the barrier separating the inactive and active conformations under different conditions. At this step, we thus hypothesize that a quantitative evaluation of the energy profiles of GHSR-1a might require a better sampling of these local motions by unbiased molecular dynamics (MD) simulations.

### Converged Markov state models capture the effect of different ligands on the energy landscape of GHSR-1a

To overcome the limitations of our metadynamics protocol and capture subtle ligand effects, we adaptively re-sampled the conformational space of the GHSR-1a systems using standard (unbiased) MD simulations upon which we constructed Markov state models (MSMs). To sample the wide range of possible configurations of each system in standard MD, we performed regular space clustering on the consolidated metadynamics trajectory data independently for all systems to ensure good initial coverage of the conformational space. For subsequent rounds of adaptive sampling, we selected new initial configurations for unbiased MD simulations by constructing an MSM and selecting microstates according to their stationary probability. This allowed us to efficiently re-explore the conformational space sampled by metadynamics simulations in an unbiased manner and to obtain a converged MSM validated using the implied timescale analysis (S8 Fig.) [34]. For each system, we projected the MSM free energy surface onto the two slowest time-lagged independent components (ICs), which were obtained by applying time-lagged independent component analysis (TICA) to the Cα-Cα distances spanning the TM region of the receptor (see Methods section for a detailed description of the molecular features used for TICA) (Fig. 2A). Because the ICs differed between systems, the resulting free energy landscapes were not directly comparable. However, IC1 encoded the same TM6 displacement motion, a well-described feature of receptor activation, in the space of the apo- and agonist-bound systems [16] (S9 Fig.). Additionally, both systems displayed two energy barriers, along IC1 (higher in energy) and IC2 (lower in energy). Both barriers, dividing the landscape into three energy basins, were lowered by the presence of the agonist. The landscape of the antagonist-bound system had only one energy barrier along IC1, while the landscape of the inverse antagonist-bound system showed a single energy well. These results are congruent with the observations made employing the metadynamics protocol, namely the restriction of the conformational space of the receptor in the presence of the antagonist and inverse agonist molecules (S10 Fig., Fig. 1D).

To enhance the understanding of the explored conformational spaces, we initially coarse-grained all MSMs exhibiting multiple energy minima (excluding the inverse agonist-bound MSM) into two metastable states validated by Chapman-Kolmogorov tests [34] (S11 Fig.). This enabled us to directly compare the state populations with biophysical studies previously conducted by our research group, which only allowed the resolution of the two most populated states [38]. For all three systems, the two obtained metastable states were separated by the energy barrier along IC1 (Fig. 2B). By comparing the RMSD of the minimal-energy microstates of each metastable state to experimental structures of GHSR-1a, we assigned the metastable states of the apo and agonist-bound systems to active and inactive receptor conformations which have been experimentally described by structures of GHSR-1a in presence of an agonist (PDB id. 7F9Y) and antagonist (PDB id. 6KO5), respectively (S12 Fig.). The projection of the experimentally resolved class A GPCR structures onto the related IC spaces unambiguously supported this state assignment with the respective metastable states overlapping well with inactive and active structures (Fig. 2B). Projections of the same set of structures onto the antagonist-bound landscape equally confirmed the sampling of the inactive state, and a new state, unknown for GHSR-1a, deviating largely from all resolved experimental structures of this receptor (Fig. 2B, S12 Fig.) but showing similar features as structures resolved for bile acids receptors [39], [40], [41] and human itch receptors [42], [43]. Here, nearly all experimental structures, independent of their conformational annotation, were projected onto the same region of the space, the TICA being unable to separate the active and inactive conformations. Lastly, the energy minimum of the inverse-agonist system corresponds to the initial inactive conformation described experimentally for GHSR-1 (PDB id. 7F83) (S12 Fig.). In this case, only one macrostate was assigned to ensure the comparability and robustness of the analysis as the obtained MSM did not resolve any stable transition on time scales above 5 microseconds (Fig. S9).

Taken together, our data showed that the apo and agonist-bound receptors explored a similar space in which active and inactive conformations were both stabilized and in equilibrium. The presence of the antagonist and the inverse agonist molecules prevented the observation of such an equilibrium and trapped the receptor in inactive conformations, the antagonist-bound system being nevertheless able to explore a new state not captured by existing structures.

To quantitatively describe the conformational equilibria, we evaluated the stationary and transition probabilities of and between the metastable states. Consistent with experimental data, we observed that agonist binding to GHSR-1a increased the stationary probability of the active state from 0.2 to 0.5 resulting in a balanced equilibrium between both states [38] (Fig. 2C). Confirming our first impression on the energy barriers of the landscapes in Fig. 2A, the mean first passage time (MFPT) towards the active state (activation process) was decreased significantly in the presence of the agonist. On the contrary, no effect was observed on the transition kinetics of the deactivation process (Fig. 2D). Together, these results show that the agonist not only stabilizes the active conformation but also acts on the transition kinetics of the activation process. In the case of the antagonist-bound system, our analysis revealed a balanced equilibrium between the inactive and the newly identified state (Fig. 2C). Transitions between these two states were significantly faster than between the inactive and active states in the apo and agonist-bound receptor (Fig. 2D). This further supports that antagonist binding to the receptor must restrict it to a new, smaller (S11) conformational space with the known inactive conformation being in fast exchange with the newly identified state.

### Refined clustering reveals intermediate states

As delineated above, dividing the various conformational landscapes into two states already allowed us to discuss the balance between active and inactive conformations (apo and agonist bound systems) or to identify a new metastable state in the case of the antagonist bound system. However, it restricted the identification of states to those separated along IC1, while we observed another energy barrier along IC2 in the apo- and agonist-bound systems (Fig. 2A). To identify the conformations governing the heterogeneous inactive energy basin of these two systems more precisely, we re-clustered the conformational landscape into three metastable states validated by Chapman-Kolmogorov tests [34] (S14 Fig.). The inactive conformations in each case were now assigned to two different metastable states effectively separated by the energy barrier along IC2 (Fig. 3A). These two states, referred to below as *intermediate 1* and *intermediate 2*, are different (Fig. 3B) and show another level of discrepancy between the apo and agonist-bound systems.

The *intermediate 1* metastable state was only sampled by the apo receptor and is reported in green in Fig. 3. It exhibits a closed conformation on the intracellular side as found in inactive conformations but a significant shift in the extracellular parts of TM5, TM6, and TM7, contributing to open the receptor (Fig. 3B). This is of particular interest as it has been previously proposed that the highly lipophilic octanoyl moiety of the ghrelin peptide, the endogenous GHSR-1a agonist, could lead to its initial partitioning in the membrane and subsequent lateral diffusion to the orthosteric binding site [44], [45]. In this context, the opening between TM5 and TM6 could create the necessary entry pathway for the ghrelin peptide, not seen in any of the GHSR-1a structures solved to date. We nevertheless argue that such a pathway has already been proposed for other GPCRs including the sphingosine-1-phosphate receptor [44] and the opsin receptor [46], [47]. Additionally, this hypothesis was supported by coarse-grained MD simulations, which we performed starting from the receptor in its active and *intermediate 1* conformations in the presence of the solvated ghrelin peptide. In these simulations, we promoted the binding of the peptide to the receptor by employing the RMSD computed against the reference structure of the complex as a collective variable. In all the obtained trajectories, we observed a spontaneous anchoring of the peptide to the membrane, mediated by interactions of its octanoyl moiety and PHE4 residue with lipids, as proposed in recent NMR studies [45]. In both cases, after partitioning into the membrane, the peptide sampled different positions around the receptor. Whereas the peptide was hindered from entering the receptor in its active conformation, in the simulations of the receptor in the *intermediate 1* conformation, the peptide diffused through the opened TM5-TM6 interface adopting its correct orientation in the orthosteric site (S14 Fig.).

To further discuss the role of the *intermediate 1* state, we quantified its transition probabilities to the known active and inactive states of GHSR-1a (Fig. 3D). *Intermediate 1* quickly transited to the inactive conformation with an MFPT of 2.8 + 0.6 µs describing a closing of the binding pathway in absence of the ligand. Note that, based on our data, we cannot discuss the rate of this transition after ligand binding. However, as the inactive conformation preferably transited towards the *intermediate 1* conformation compared to the active one (MFPT = 20 + 3 µs versus MFPT = 83 + 23 µs), we hypothesize that the transition via this state constitutes an alternative activation path. Indeed, although the transition from *intermediate* 1 to the active state is relatively slow without the ligand bound (MFPT = 85 ± 22 µs), we cannot rule out the possibility that the agonist may accelerate this process.

The additional metastable state of the agonist-bound system (*intermediate 2*) is reported in light blue in Fig. 3. After visual inspection, we concluded that this state described the same conformation as the above-identified new state of the antagonist-bound GHSR-1a system. The conformer was also sampled by the apo system, note, however, that in this case, the respective microstates were not assigned to a distinct metastable state (Fig. 3A). *Intermediate 2* presented an open conformation with a TM1-TM6 distance of 26.5 + 2.7 Å comparable to the Gq-bound cryo-EM structure of GHSR-1a (PDB id. 7F9Y, d_TM1-TM6_ = 26.2 Å). In contrast, it exhibited an outward shift of the intracellular region of TM5 and a displacement of the intracellular part of TM6 towards TM5 (Fig. 3C). The resulting kink in TM6 has been extensively described for other class A GPCR including bile acids receptors [39], [40], [41] and human itch receptors [42], [43], with all structures being G-protein-coupled. As, in these studies, the kink mainly stabilized by the interaction of T^3.36^ and F^6.51^ (corresponding to T127 and F279 in GHSR-1a) was described to be crucial in G-protein coupling and receptor activation, we hypothesize it to play a similar role in the *intermediate 2* conformation. Indeed, the fast and balanced equilibrium with the inactive conformation, with an MFPT in the order of ∼10 µs for the agonist- and antagonist-bound systems pointed to *intermediate 2* as a preferred gateway to the active state. However, since in the presence of the agonist, the transition between the *intermediate 2* and the active conformations occurred at the same rate as the direct inactive → active transition (MFPT = 20 + 3 µs), it is difficult to conclude a predominant activation pathway. We can only hypothesize here that the alternative activation pathway might be favored in the presence of an intracellular protein partner binding to this intermediate state.

### Agonist facilitates GHSR-1a activation by constraining the transition path

As outlined above, the direct transition between the inactive state and the active state was four times faster when the agonist was bound to the receptor compared to the receptor in its apo form. To elucidate the molecular details of how the agonist facilitates this transition, we used transition path theory [48], [49] to sample the ten most probable paths from the inactive to the active state for both systems (Fig. 4A). In both cases, the trajectories followed IC1 without visiting *intermediate 1* or *intermediate 2*. This is not surprising, as we discussed above, how ligand and G-protein binding to *intermediate 1* and *intermediate 2* respectively, would be necessary to favor these two alternative pathways. The projection of the transition paths onto the TICA space suggested that the conformational space explored during activation was restricted when the agonist was bound to the receptor (Fig. 4A). This was confirmed by the root mean square fluctuation (RMSF) profiles of the microstates sampled during the transition (Fig. 4B). The agonist restricted the flexibility of several transmembrane helices, most prominently of TM1, TM6, and TM7. To explore the modulation of the transition path and potentially the modulation of the allosteric network caused by agonist binding in greater detail we used force distribution analysis (FDA). Indeed, forces offer a more precise measure, revealing low-amplitude but functionally critical motions that trigger allosteric communication and subsequent conformational modifications. We retrieved the pairwise forces within the receptor when it was in the agonist-bound state, relative to the force in the apo state at the start of the transition path to capture subtle reorganizations directly linked with the presence of the ligand. (Fig. 4C). The pairwise forces significantly changed for several residue pairs in the intra- and extracellular loops which we attributed to their high flexibility. Interestingly, in the GHSR-1a core region, only a few residue pairs showed significant changes in their pairwise forces, indicating a large redistribution of mechanical stress among them. These residues included several so-called “microswitches” which are hypothesized to govern the allosteric communication between the agonist-binding pocket and the intracellular region [6], [50], [51]. Namely, these included V131^3.40^, F272^6.44^, and W276^6.48^ known as transmission switch residues [14], Y323^7.53^ of the NPxxY-motif [16] as well as D89^2.50^, T127^3.36^, and F312^7.42^, which are involved in sodium binding, a known allosteric modulator for GHSR-1a (Fig. 4C) [52]. Taken together this indicates that the agonist facilitates the receptor activation path by reducing the entropy along the transition path restricting the main force transmission from the extracellular to the intracellular side to a few key residues before the actual conformational transition. When considering the full transition path, it is clear that the receptor underwent highly global motions that drastically altered the interaction network within the receptor (S16B Fig.). Interactions between TM5 and TM6 as well as between the C-terminus part of TM7 with TM1 and TM2 were drastically modified along the activation pathway. Interestingly, TM3 had a highly central role in the allosteric communication, while TM4 seemed to not have any significant impact on the activation motion of GHSR-1a, these observations were in line with a previous study based on class A GPCR structures [53].

**Figure 4:**
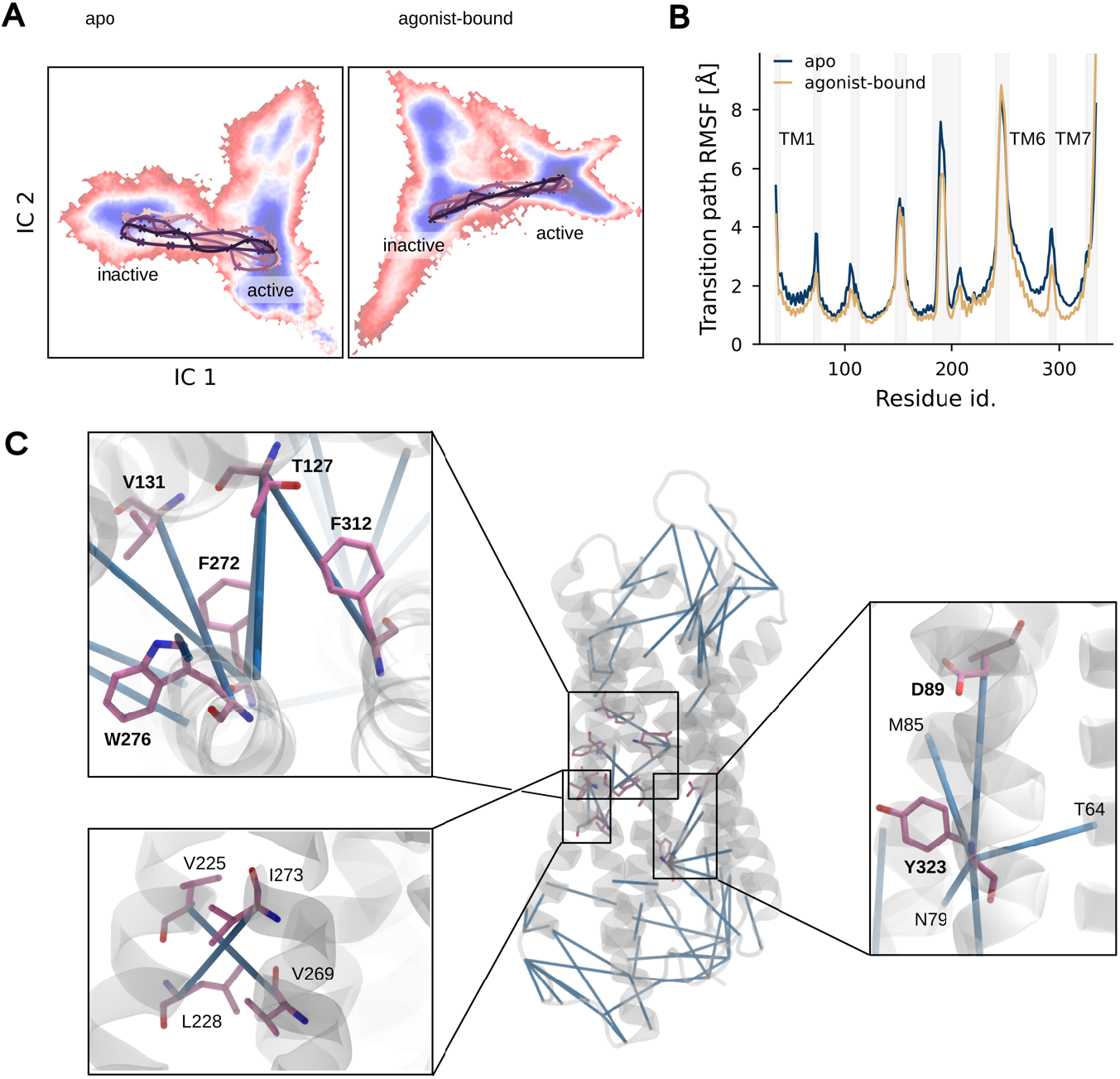
Mechanistics of the agonist effect on GHSR-1a activation process. A: MSM free energy landscape of the apo (left) and agonist-bound (right) GHSR-1a with the n=10 most likely transition paths between the inactive and active states. The order is indicated with a color gradient from purple to light pink. B: Root mean square fluctuation (RMSF) of all microstates along the transition paths. C: Shown is the change in pairwise force ΔF_ij_ = 〈F_ij_(agonist-bound)〉− 〈F_ij_(apo)〉for the pair of residues (i, j). ΔF is depicted as blue lines connecting the residues (i, j). Only pairs displaying a statistically significant change are shown: score z > 0.75 and |ΔF| > 50 pN. Residues displaying significant changes are highlighted in pink. Labels of residues that are known microswitches are highlighted in bold.

## DISCUSSION

In this work by using metadynamics simulations and Markov state modeling, we shed light on the conformational landscape of GHSR-1a, a member of the class A GPCR family, and its modulation by different classes of ligands (agonist, antagonist, and inverse-agonist). Our rational approach to obtaining CVs for bias-exchange metadynamics simulations from PCA of experimental class A GPCR structures not only proved successful for our systems but is inherently applicable to other proteins of this family. Our results demonstrate the effectiveness of the chosen CVs in exploring active and inactive conformations of GHSR-1a when in its apo form or bound to an agonist (Fig. 1). Furthermore, the metadynamics simulations qualitatively captured the effect of different ligands on the internal dynamics of GHSR-1a. This is a remarkable result as finding robust CVs for biased MD simulations of GPCRs has been a key challenge in the field [7], [8], [9], [10], [11], [12]. Since our collective variables are transferable to other class A GPCRs, this approach may be of particular interest for a relatively rapid assessment of the effects of different ligands on the equilibria of other GPCRs. Note, however, that our metadynamics protocol failed to quantitatively detect the effect of the agonist on the energy landscape of the receptor, despite its known role in stabilizing the active state. While for the apo and agonist-bound system, we captured the inactive and active states separated by an energy barrier, high variability between replicas prevented us from identifying any statistically significant differences in the resulting profiles. This could be due to the chosen CVs, which enhance the sampling of highly collective motions, to the detriment of an extensive sampling of subtle local rearrangements as for example side-chain reorientations that are equally important for receptor activation [6], [50], [51].

To describe more accurately the energy landscape of the receptor, we have constructed MSMs for GHSR-1a in its apo state and bound to pharmacologically distinct ligands, based on unbiased MD simulations with a range of diverse snapshots from our metadynamics simulations as initial conformations (Fig. 2, Fig. 3). We showed that starting from different conformations of the receptor, sampled in metadynamics, allowed an efficient convergence of the MSMs. The obtained models show high agreement with a fluorescence spectroscopy study previously published by our group [38]. Our computational study not only resolves the conformational landscape of GHSR-1a in greater detail but also provides structural and kinetic information in addition to the population ratio of each metastable state. In total, our models captured five metastable GHSR-1a conformations varying in their population ratio dependent on the bound ligand, namely the active, antagonist-inactive (corresponding to PDB id 6KO5), inverseagonist-inactive (corresponding to the PDB if 7F83), *intermediate 1* and *intermediate 2* states.

In its apo form, the receptor mostly populated the antagonist-inactive state but also explored the active state, an *intermediate 1* state exposing a ligand-binding channel (Fig. 3B), and an *intermediate 2* state which we hypothesize to allow for precoupling of intracellular protein partners (Fig. 3C). Even though the intermediate states were not resolved in the fluorescence spectroscopy study as only the two main conformers could be revealed with the experimental setup, the population of the active state in absence of the agonist is in agreement with the uncommonly high constitutive activity of GHSR-1a [54], [55], [56]. Interestingly, the apo state could populate more conformers than any other system tested in this study, showing its high plasticity and conformational variability. The flexibility of the apo form can be inferred from the lack of structural data of GHSR-1a in the absence of molecular partners. Indeed, its conformational heterogeneity makes it difficult to stabilize the receptor as it constantly transits between inactive, active, or intermediate states. Similarly, the two intermediate states we observed were not solved experimentally for GHSR-1a, although prominent features of *intermediate 2* states were suggested for other GPCRs [39], [40], [41] [42], [43]. Their lifetimes and relatively fast equilibrium with the inactive conformation are challenging to catch experimentally. The design of conformation-specific binders that could stabilize these intermediate states is under consideration, as they could be interesting pharmacological tools to study GHSR-1a.

While the apo system showed the highest plasticity, the inverse agonist-bound receptor was characterized by a single, inverse agonist-inactive conformation [16], distinct from any metastable states explored by the other systems. Interestingly, the presence of the inverse agonist drastically restricted conformational sampling around the minimal energy structure resolved experimentally. The effect of the inverse agonist is in full agreement with previous experimental observations indicating a complete G-protein release from GHSR-1a in the presence of such a ligand and the stabilization of a conformational state that does not allow for G-protein precoupling [37], [38]. Interestingly, the inverse-agonist state was not sampled in the apo state suggesting that the inverse agonist promotes a super-inactive which is not sampled in the absence of the ligand.

To a lesser extent, the antagonist equally restricted the conformational plasticity of the receptor. Although the receptor was able to sample the antagonist-inactive and *intermediate 2* states, it did not explore the active state anymore. The drastic modulation of the GHSR-1a conformational landscape by this ligand contradicts the model of neutral antagonists that competitively inhibit agonist binding without affecting the GPCR equilibrium and consequently the constitutive activity of GHSR-1a [57], [58]. Our observations are however in line with the multi-state GPCR model where the presence of a neutral antagonist is being challenged [59], [60]. Additionally, it is important to note that intracellular partners were missing in our simulation setup which could modify the conformational landscape of the antagonist-bound receptor. In the presence of the G protein, we hypothesize comparable macroscopic activity of GHSR-1a in its apo state or bound to so-called antagonists in the presence of the G-protein emerging from the active or *intermediate 2* states, respectively [61].

Not surprisingly, the agonist bound to the receptor shifted the conformational equilibrium toward the active state, although the receptor could still populate the antagonist-inactive and *intermediate 2* states to a lesser extent. The fact that the apo and agonist-bound systems could explore both antagonist-inactive and active states, allowed us to study the differences in the state transitions depending on the ligand. In this context, we hypothesize that the transition characteristics of the apo and agonist-bound system correspond to the basal and ligand-induced activation of the receptor, respectively. Based on the transition path analysis performed on both MSMs a direct transition between both metastable states seemed to be the most probable path (Fig. 4A). The conformations explored along this path exhibited a smaller variability when the ligand was bound to the receptor, leading to a narrower activation path (Fig. 4B), which may explain the higher activation rate (lower MFPT) of the agonist-bound system (Fig. 3D). FDA performed on the conformations of at the start of the activation paths confirmed the increase of mechanical stress on a few key residues, widely described as microswitches in the literature, in the receptor core when the agonist is bound to the receptor [6], [50], [51] (Fig. 4C). Ghrelin binding in this context then appeared not only to stabilize the active state of the receptor but also to reduce the entropy of the corresponding transition. Our results suggest that this effect could be explained by the exertion of forces on residues belonging to the so-called “microswitches”. Note here, that this affects “microswitches” throughout the receptor core in a simultaneous manner further substantiating global motions as the driver of receptor activation.

Despite the insights our MSMs provided on the ligand effect on receptor activation, they lack a comprehensive view including the effect of intracellular partners (e.g. G-proteins, β-arrestins or kinases), which are known to have a strong effect on the conformational landscape of GHSR-1a [38]. For example, even though the direct transition between the inactive and active state was predicted to be the most probable when the agonist is bound, the transition inactive → *intermediate 2* is significantly faster than the transition inactive → active (Fig. 3D). As we hypothesize the *intermediate 2* state to allow for intracellular protein partner pre-coupling, it seems probable that the rate of the transition *intermediate 2* → active might decrease in presence of these partners. However, due to the high computational cost of the unbiased MD simulations required to construct the MSMs the application of the proposed protocol to bigger systems such as GHSR-1a:G-protein complexes is outside the scope of this work. The use of coarse-grained force fields such as MARTINI 3 [62], [63], would allow us to decrease its computational cost and enable us to equally study the landscape modulation by intracellular protein partners. However, to date, constraints (i.e. Go-bonds) used to maintain the tertiary structure of the receptor prevent any large-scale motions. Modification of these constraints to improve sampling is under consideration for future work. In that perspective, our energy landscapes could serve as objective functions for optimization to enable the coarse-grained force field to reproduce realistic motions that respect the thermodynamic properties of GHSR-1a.

## ASSOCIATED CONTENT

### Supporting Information

The Supporting Information is available free of charge on the ACS Publications website.

Molecular dynamics simulation details, technical details of data analysis technical details, convergence of metadynamics simulations, construction and validation of Markov state models, technical detail for ghrelin binding simulations, force distribution analysis (PDF)

## AUTHOR INFORMATION

### Author Contributions

R.R., N.F., and M.L. designed the research. R.R. performed and analyzed the MD simulations. R.R., N.F., and M.L. wrote, reviewed, and edited the paper.

## ACKNOWLEDGMENT

We thank the CNRS, the Université de Montpellier, the Ecole Nationale Supérieure de Chimie de Montpellier (ENSCM), and the Agence Nationale de la Recherche (Projet-ANR-22-CE44-0042) for their financial support. This work was granted access to the HPC resources of IDRIS and CINES under the allocations A0140714133 and A0160715115 made by GENCI. We thank Alessandro Barducci, Patrick Fuchs, Paula Milán Rodríguez, and Jean-Louis Banères for helpful discussions.

## Supplementary Materials

## Methods

### Systems setup

The initial models of the ghrelin-bound and apo GHSR-1a were built from the available cryo-EM structure of this receptor bound to its endogenous agonist peptide and to the G-protein Gq (PDB id. 7F9Y) [1]. For both systems, the G-protein was removed. In the case of the apo-GHSR, the ligand was also deleted. The antagonist-bound and inverse agonist-bound initial models of the receptor were obtained from existing X-ray structures of GHSR-1a bound to synthetic small molecules (PDB ids. 6KO5, 7F83) after the removal of lipids and stabilizing proteins [2], [3]. Missing residues and loops were completed using Modeller 10.4 [4]. The models were embedded in symmetric lipid bilayers composed of POPC using the CHARMM-GUI web server [5], [6], [7]. The proteins were positioned within the membrane according to the results from the “Orientation of Proteins in Membranes” (OMP) server [8]. The canonical disulfide bond of class-A GPCRs was added between C116 and C198 of the receptor. A CHARMM patch was created to covalently link an octanoate moiety to the hydroxyl group of SER3 of ghrelin. The systems were solvated with water and 150 mM (Na+, Cl-).

### Simulation algorithms and parameters

All simulations described in this work were carried out using GROMACS 2021.5 [9], the CHARMM36m force field [10], [11], and the CHARMM-modified TIP3 water model [12]. All hydrogen bonds were constrained using the LINCS algorithm [13]. An integration time step of 2 fs was used unless otherwise indicated. Long-range electrostatic interactions were treated using the Particle Mesh Ewald (PME) [14] technique and the Verlet cut-off scheme whereby neighbor lists with a cut-off of 1.2 nm were updated every 10 steps. Non-bonded interactions such as electrostatic and van-der-Waals interactions were truncated at 1.2 nm. The energies of all systems constructed in this study were minimized using the steepest descent method with a tolerance of 1000 kJ·mol-1·nm-1 and a maximum of 5000 steps. The minimization was followed by NVT and NPT equilibration steps according to the CHARMM-GUI protocol where heavy atom position restraints were gradually removed. Subsequently, the system was equilibrated in the NPT ensemble for 100 ns with only backbone atoms being weakly restrained with a force constant of 50 kJ·mol-1·nm-2. For all equilibrium simulations, the temperature was maintained at 303.15 K through the use of the Berendsen thermostat [15], coupling protein, membrane, and the rest of the system separately to the thermostat, and using a coupling time constant of 1 ps. For production, a velocity-rescaling thermostat [16] was used with the same parameters. The pressure was kept constant at 1 bar employing a semi-isotropic Berendsen barostat [15] for equilibration and a semi-isotropic Parrinello-Rahman barostat [17] in production (coupling time constant of 5 ps).

### Coarse-grained simulations of ghrelin binding to GHSR-1a

Coarse-grained (CG) MD simulations were performed using the MARTINI 3 force field [18]. The active and intermediate 1 states obtained in our simulations of the apo GHSR-1a as well as the octanoylated ghrelin peptide obtained from PDB id. 7F9Y were converted to coarse-grained resolution using Martinize2 [19]. An elastic network was employed (dcut-off = 0.9 nm, fc = 700 kJ·mol-1·nm-2) to preserve the tertiary structure of GHSR-1a. The insane.py script [20] was used to embed the coarse-grained receptor in a lipid bilayer and to solvate the final system. We used a membrane composition comparable to our previous studies of GHSR-1a (48 % POPC, 27 % POPG, 5 % PI(4,5)P2, 20 % Cholesterol) [21]. The ghrelin peptide was placed at an initial position of 3 nm above the membrane surface, removing overlapping water beads. The global charges of the two systems were neutralized by adding Na+ and Cl-ions to reach a concentration of 0.15 M. The systems were energy minimized and equilibrated for 50 ns (for more details see: Simulation algorithms and parameters). All coarse-grained simulations were carried out with a 20 fs timestep. For production, well-tempered metadynamics simulations [22] were carried out using the PLUMED 2.8.1 plug-in [23], [24] incorporated into GROMACS [25]. During the 500ns-long production simulations, position restraints were maintained on all backbone beads of the receptor excluding those located in loops to maintain the global shape of the receptor’s orthosteric site whereas side chains of all residues were free to move. The RMSD computed against the reference structure of the ghrelin-bound GHSR-1a (PDB id. 7F9Y, coarse-grained with Martinize2) was used as a collective variable. Gaussian hills with an initial height of 1.0 kJ·mol−1 and width of 0.1 Å were deposited every 10 ps with.

### Transition path analysis

We used transition path theory (TPT) implemented in the Deeptime library [26] to compute the most probable transition pathways from the inactive to the active metastable state for the apo and agonist-bound GHSR-1a system (Fig. 4A). Based on the MSM transition matrix we calculated forward and backward committor probabilities, quantifying the likelihood of reaching the target microstate before returning to the starting state. By decomposing the resulting flux matrix, we identified the 10 highest-capacity paths that capture the most significant transitions from the starting (= lowest-energy microstate of the inactive state) to the target microstate (= lowest-energy microstate of the active state). For each system, we computed the root-mean-square fluctuations (RMSF) on the Cα atoms of all microstates visited along the 10 most probable paths using the MDAnalysis.analysis.rms module (Fig. 4B) [27].

### Force distribution analysis

The pair-wise force F_ij_ between the pair of residues (i, j) was obtained using the force distribution analysis (FDA) tool implemented in GROMACS (version 2.11) [28]. The pair-wise force difference ΔF_ij_ = 〈F_ij_(agonist-bound)〉− 〈F_ij_(apo)〉was calculated with “agonist-bound” and “apo” indicating the respective GHSR-1a system. 〈〉denotes an ensemble average. In practice, averages of 100 frames assigned to the lowest-energy microstate of the metastable inactive state were computed separately for both systems. The following z score function was evaluated z = ΔF_ij_ / sqrt[σ^2^(F_ij_(agonist-bound)) + σ^2^(F_ij_(apo))] to give an idea of the statistical significance of the change, with σ denoting the standard deviation over the 100 frames. Pair-wise force differences that exceeded 50 pN at a z threshold of 0.75 were shown (Fig. 4C, S16A Fig.).

## Supplementary Data

**S2 Figure:**
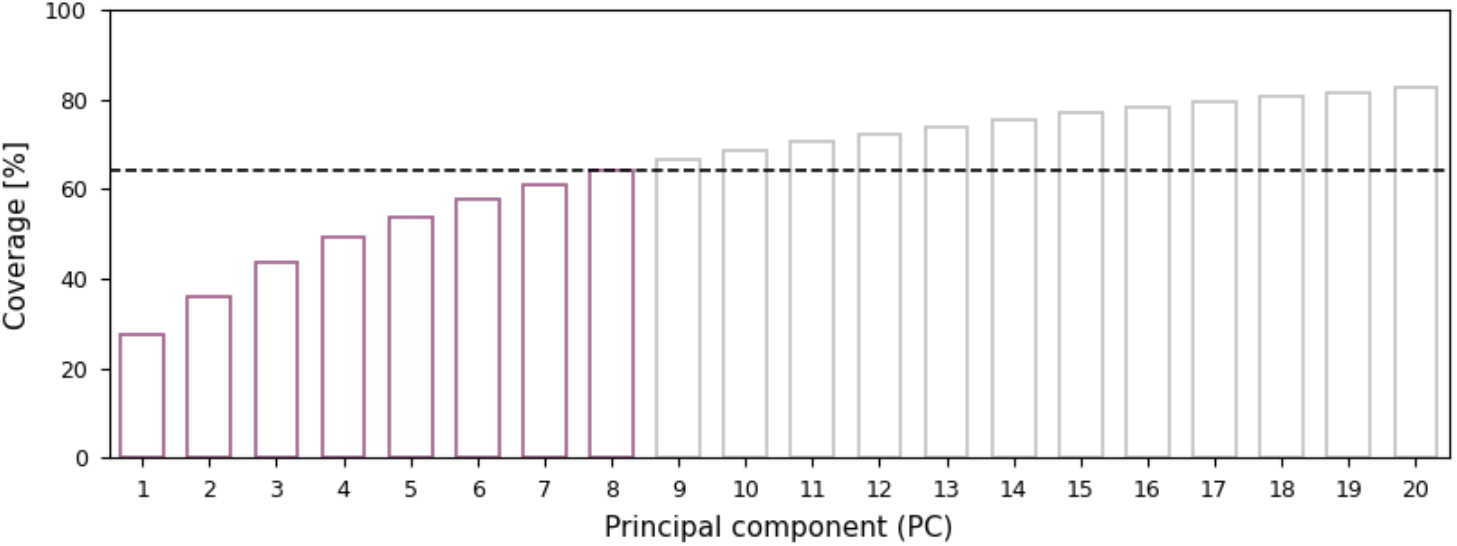
Percentage of the cumulative variance described by PCA as a function of the respective PCs. The dashed line indicates the value for the eight PCs (64 %) that were considered as collective variables for the metadynamics simulations.

**S3 Figure:**
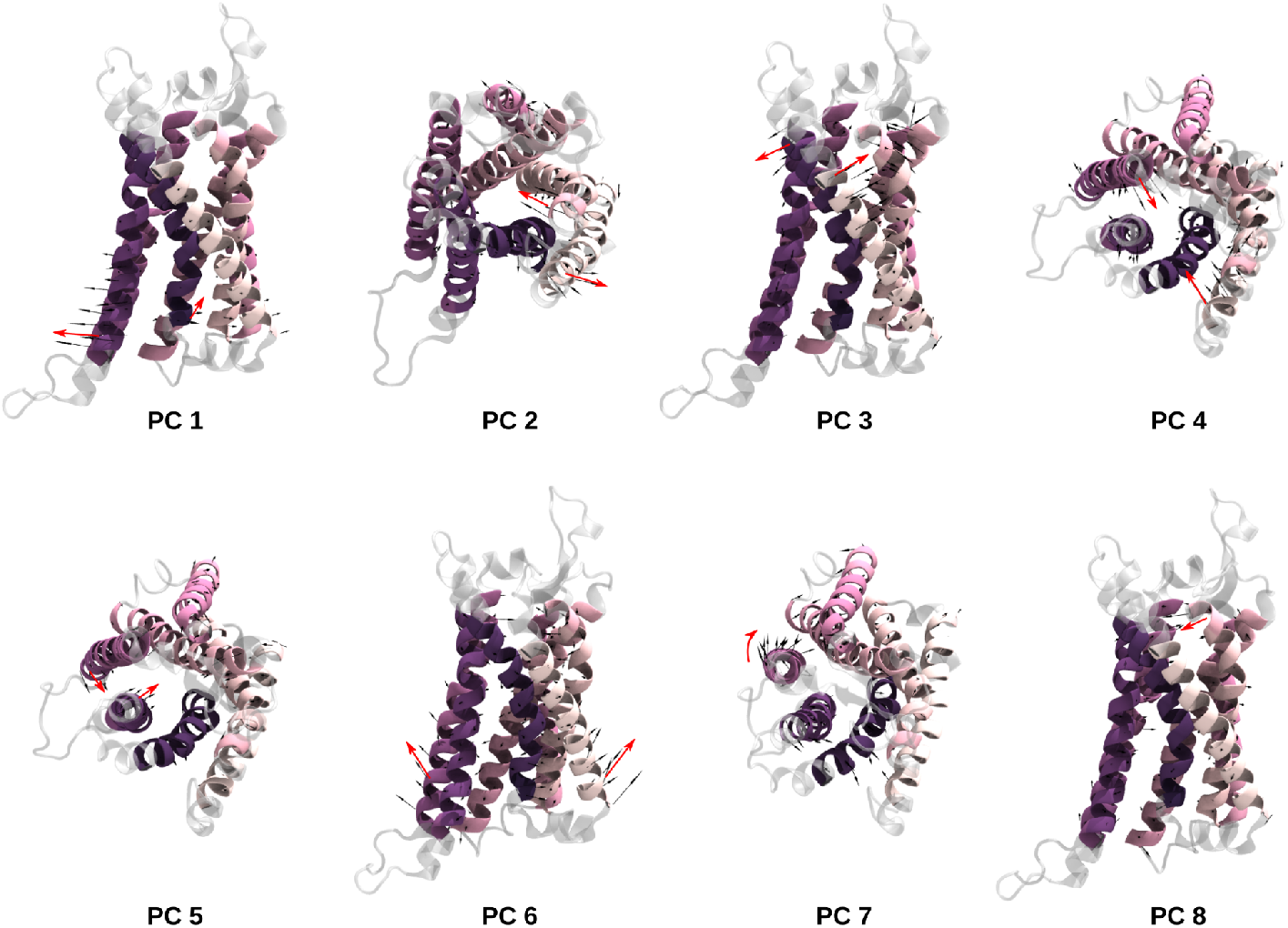
Collective motions along the first eight principal components. GHSR-1a is shown in cartoon representation. The residues of the membrane-spanning region considered in PCA are colored by index (pink to purple); the remaining residues are shown in gray. The black arrows indicate the collective motion along the respective PC, with the arrow length corresponding to the amplitude of the vector. The red arrows highlight the largest rearrangements.

**S4 Figure:**
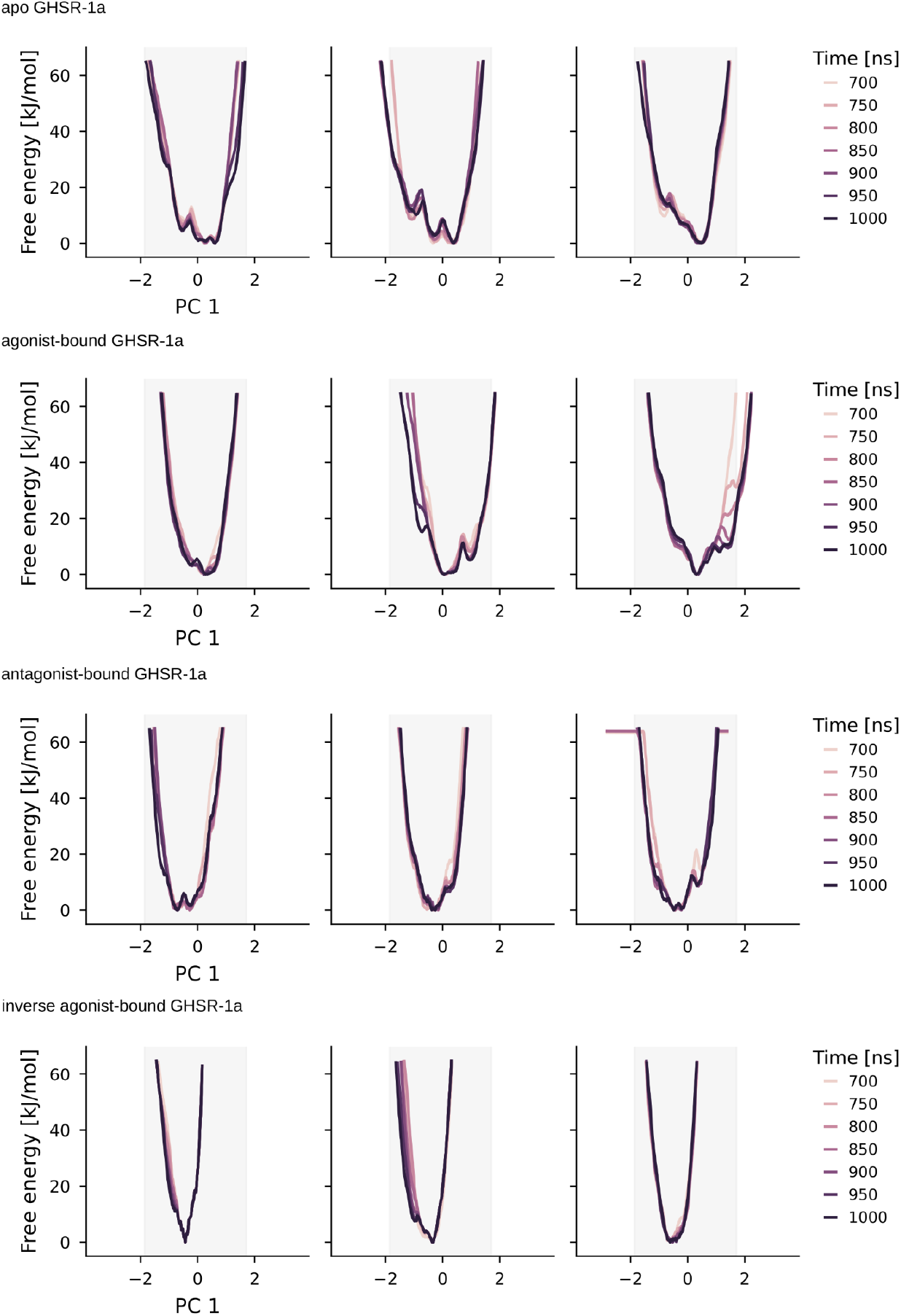
Time dependence of calculated free energy profiles obtained from metadynamics simulations. Traces as a function of PC1 are visualized for all n=3 replicas of apo, agonist-bound, antagonist ghrelin-bound, and inverse agonist-bound systems. The color gradient from light to dark indicates the progress of the simulations. The profiles remain unchanged over the last 300 ns, indicating that the simulations have converged.

**S5 Figure:**
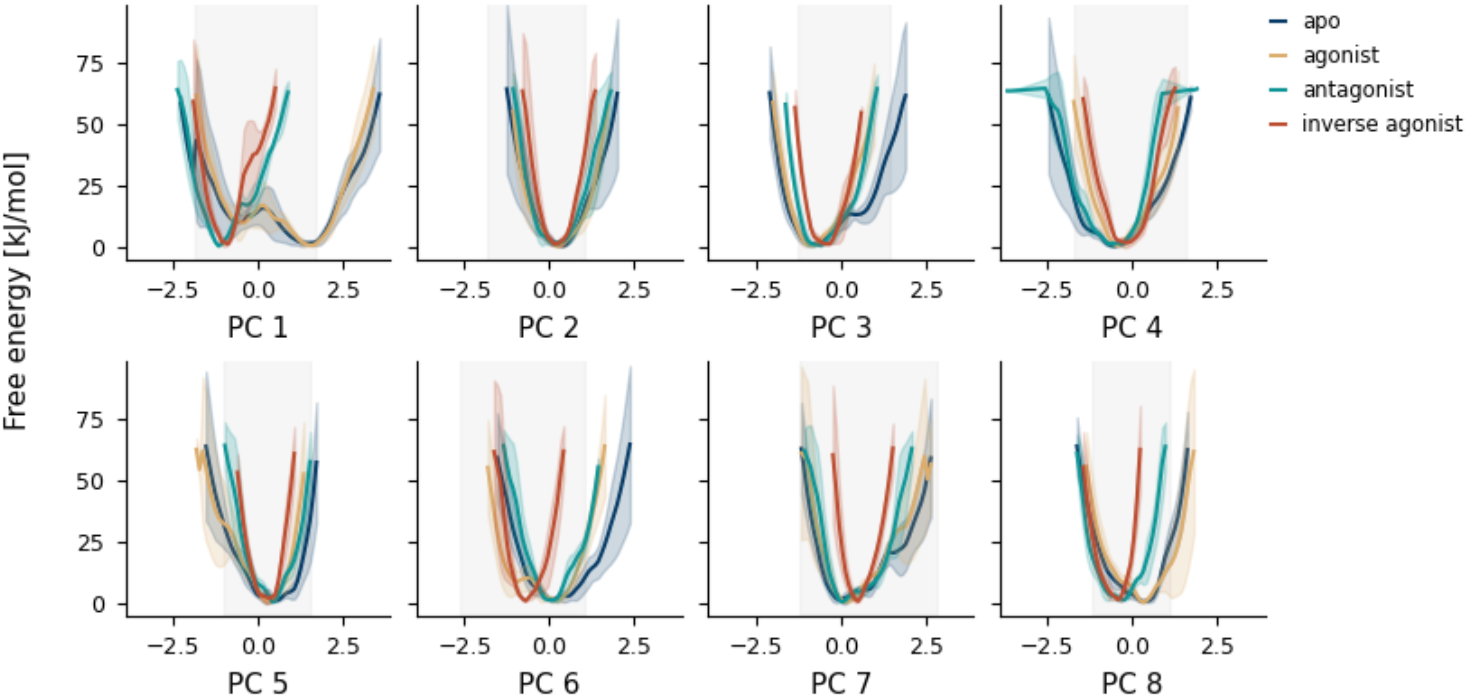
Free energy profiles obtained from metadynamics simulations. Average ± standard deviation of free energy profiles (n=3 independent replicas) as a function of PC1 to PC8 for the apo-(green), ghrelin-bound, (yellow), antagonist-bound (green), and inverse agonist system (red). The area in which the projections of the experimental structures are found is depicted in light grey.

**S6 Figure:**
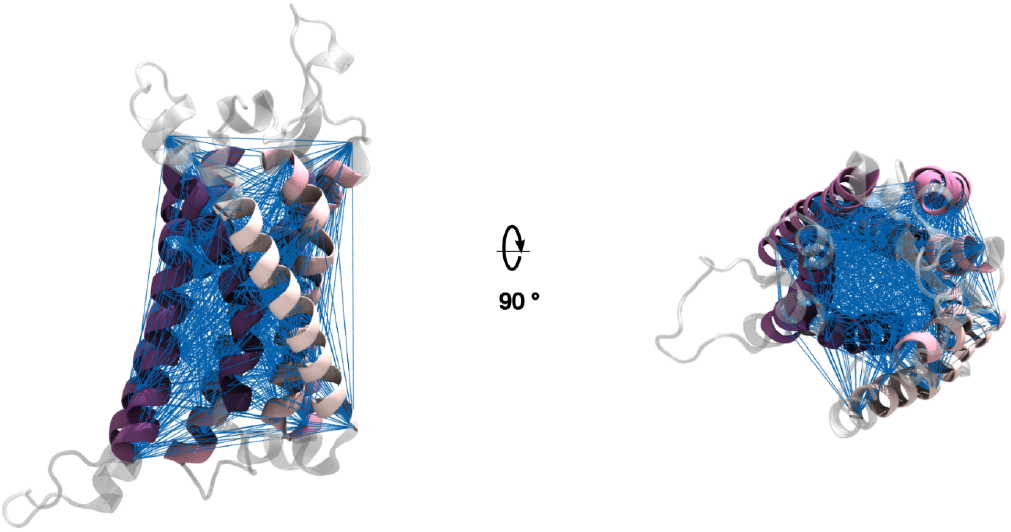
Cα distance features used as TICA input. GHSR-1a is shown in cartoon representation. The residues of the membrane-spanning region considered in PCA are colored by index (pink to purple); the remaining residues are shown in gray. The distances between the Cα atoms that we used for TICA and subsequent MSM construction are depicted as blue lines.

**S7 Figure:**
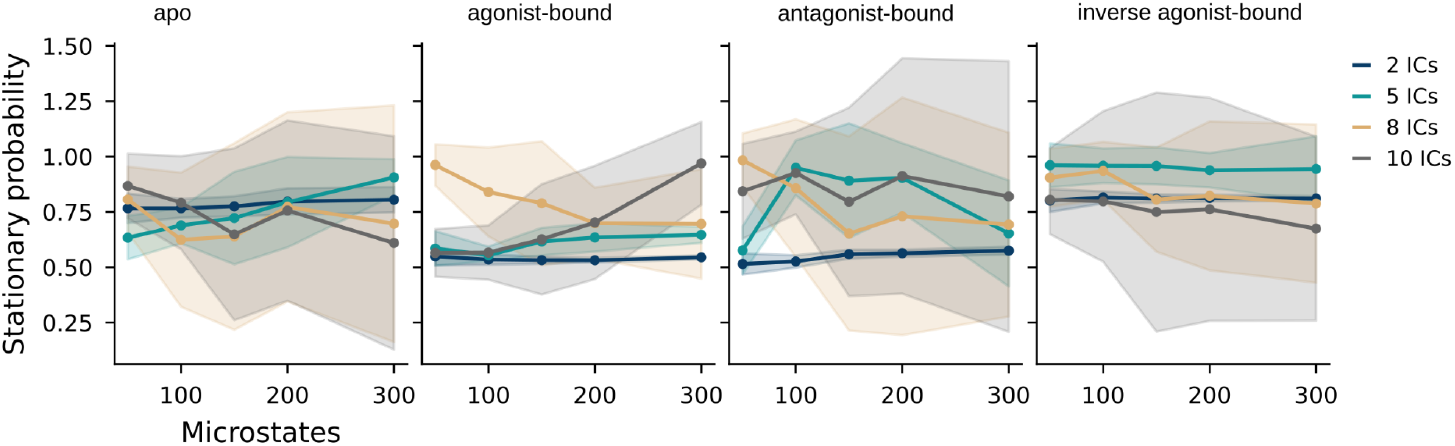
Optimization of MSM hyperparameters. The VAMP2 scores are calculated at varying numbers of microstates and TICA dimensions (ICs) for the apo, agonist-bound, antagonist-bound, and inverse agonist-bound systems. The mean values from five-fold cross-validation are plotted as dots and standard deviations as shaded regions.

**S8 Figure:**
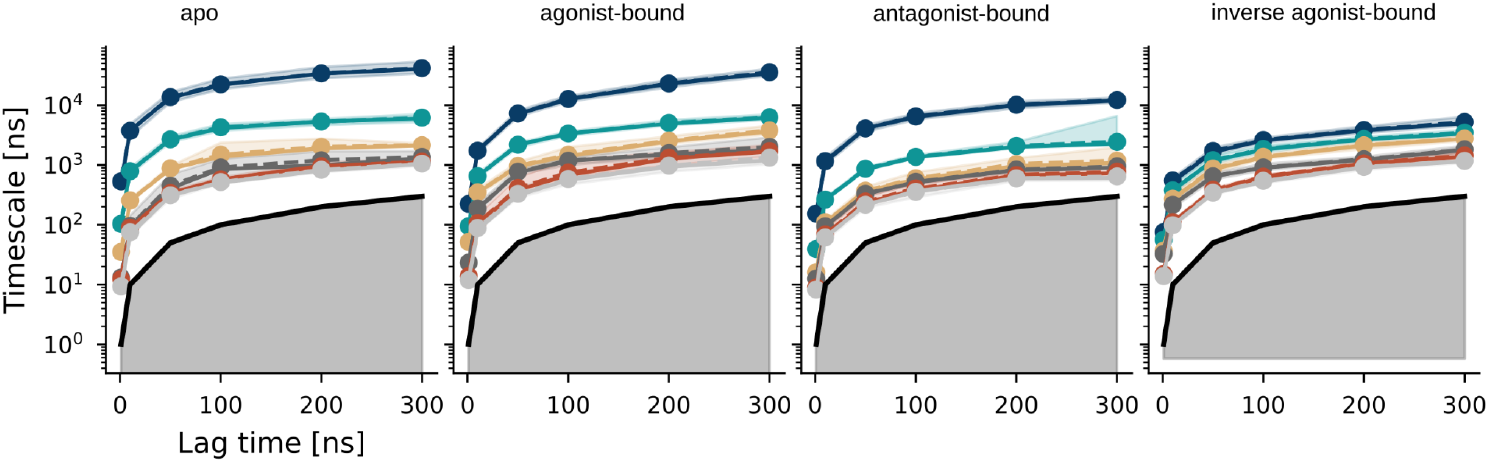
Implied timescale analysis to identify a memoryless Markovian time. The top six eigenvalues of the transition probability matrix for the apo, agonist-bound, antagonist-bound, and inverse agonist-bound systems are calculated at varying lag times. The solid lines correspond to the implied time scales of maximum likelihood MSMs. The 95% confidence intervals, quantified based on a Bayesian scheme, are depicted as shaded regions. The sample means are given by dashed lines. The black solid curve indicates the time resolution limit of the estimated Markov model.

**S9 Figure:**
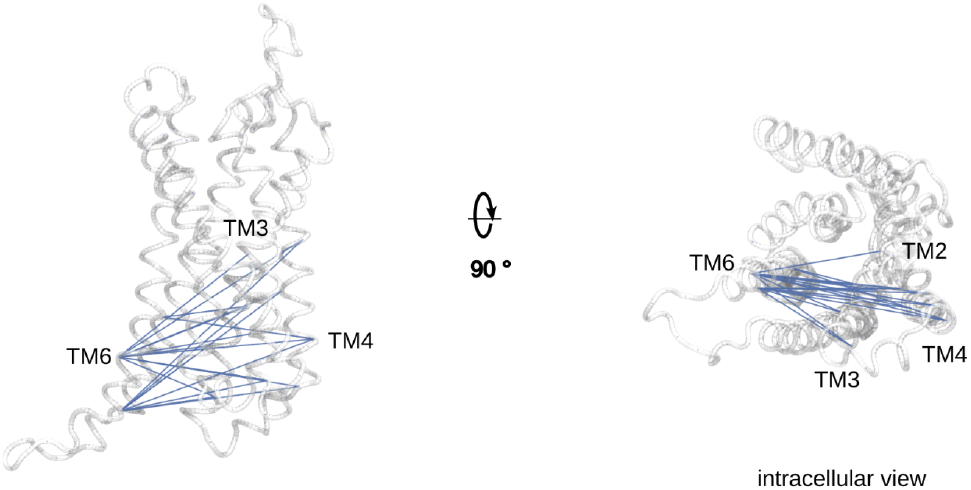
Side and intracellular view of GHSR-1a represented as ribbons with the 20 Cα-distances exhibiting the largest correlation with IC 1 highlighted in blue.

**S10 Figure:**
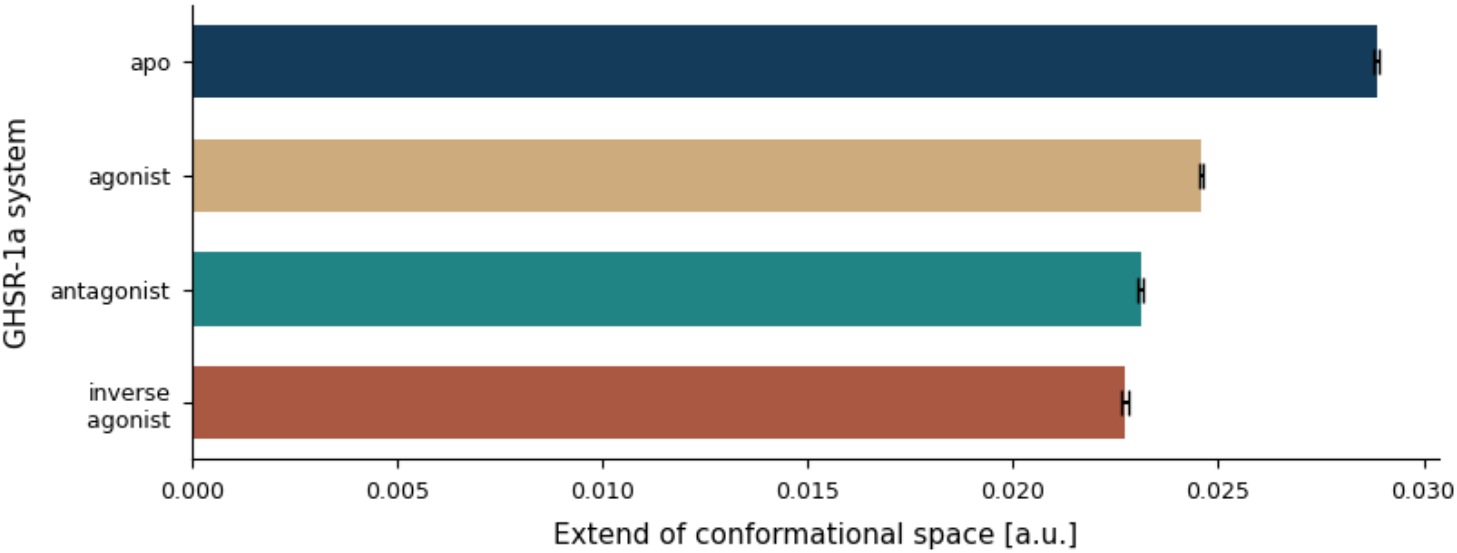
Extend of the conformational space explored by each system during unbiased MD simulations on the basis of which the MSMs were constructed. We calculated the Pearson correlation coefficient of the Cα-distance features between each trajectory frame used for MSM construction and a reference (PDB id. 7F9Y). We calculated the extent of the correlation value distribution, excluding outliers whereby the values presented in the barplot indicate the mean ± standard deviation obtained from bootstrapping using 1000 samples.

**S11 Figure:**
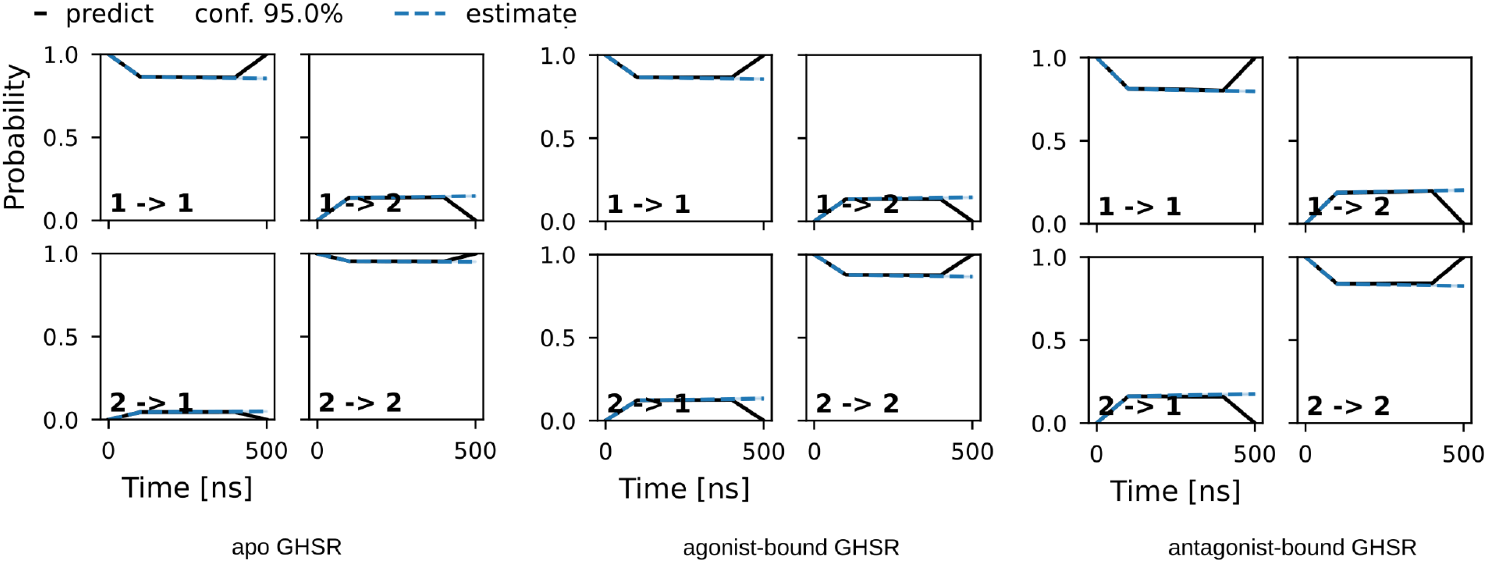
Chapman-Kolmogorov test to assess the kinetic self-consistency of the MSM. The tests for the apo, agonist-bound, and antagonist-bound systems indicate that the model predictions (black line) at long time scales are consistent with the corresponding estimates (blue line). Estimates are shown with 95% confidence intervals.

**S12 Figure:**
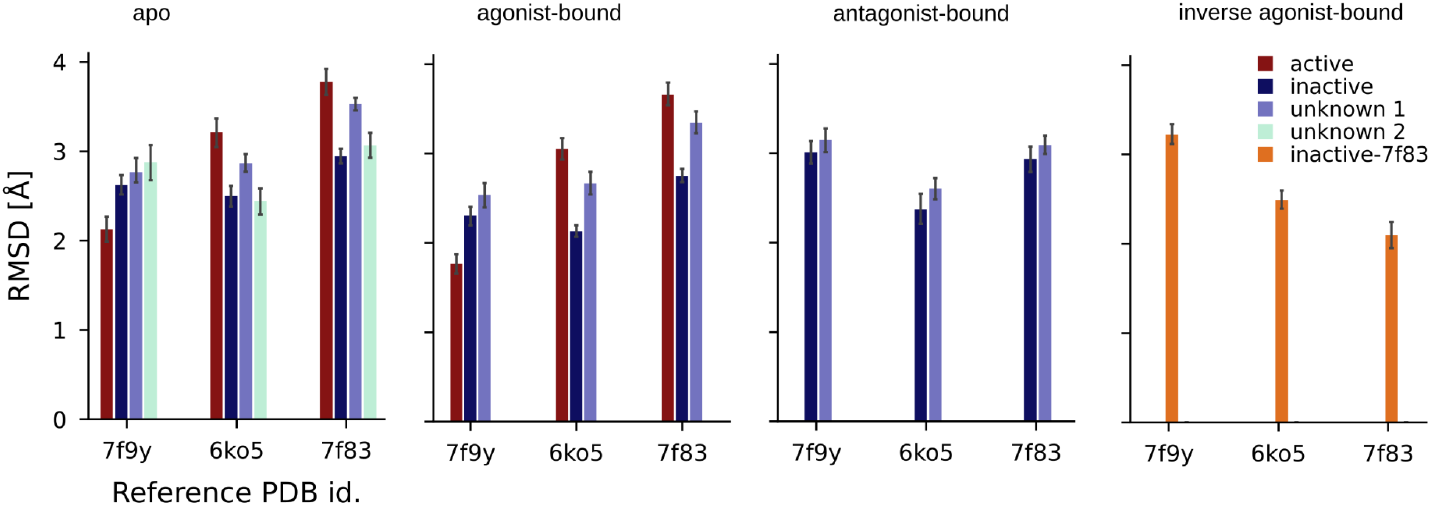
RMSD (average ± standard deviation) of the lowest energy models of each macrostate of all GHSR-1a systems with respect to the experimental structures of the agonist-(PDB id. 7F9Y) antagonist-(PDB id. 6KO5) and inverse agonist-bound receptor (PDB id. 7F83).

**S13 Figure:**
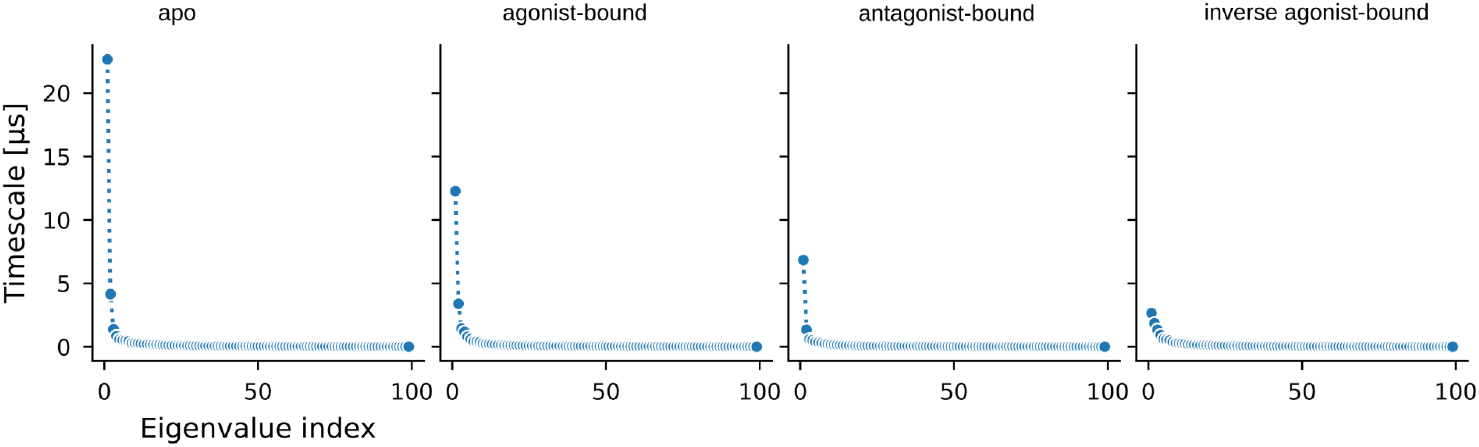
Spectral analysis of the eigenvalues of the apo, agonist-bound, antagonist-bound, and inverse agonist-bound system to identify the number of PCCA+ clusters.

**S14 Figure:**
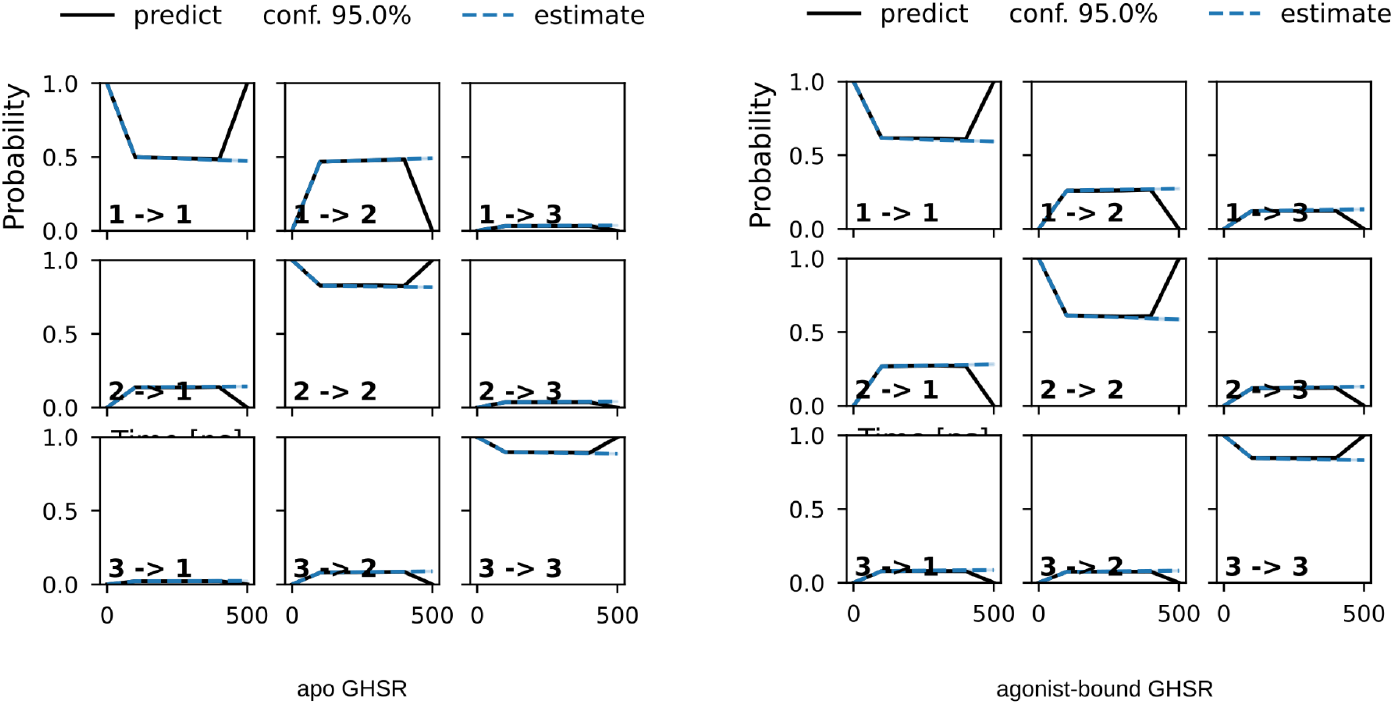
Chapman-Kolmogorov test to assess the kinetic self-consistency of the MSM. The tests for the apo, and agonist-bound systems indicate that the model predictions (black line) at long time scales are consistent with the corresponding estimates (blue line). Estimates are shown with 95% confidence intervals.

**S15 Figure:**
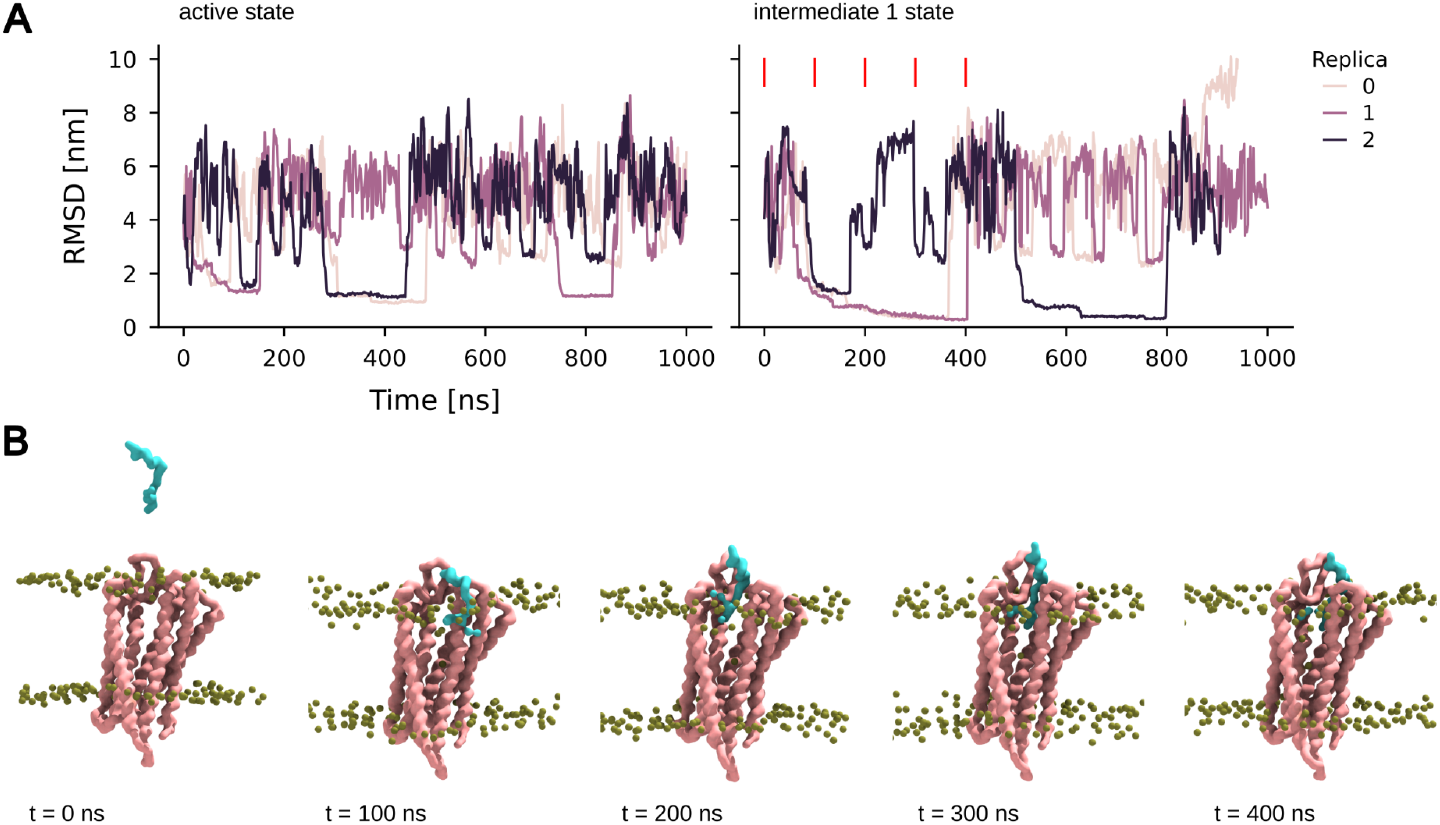
Ghrelin binding to GHSR-1a in intermediate 1 state in CG metadynamics simulations. A: Time traces of the RMSD computed ghrelin peptide backbone bead compared to the coarse-grained reference structure of the ghrelin-bound receptor (PDB id. 7F9Y) along the metadynamics simulations of GHSR-1a in its active (left) or intermediate 1 conformer (right). Different colors correspond to different replicas (n=3 per system). The red lines indicate the time points of the snapshots in B taken from replica 1 of the intermediate 1 simulations. B: Snapshot along the CG metadynamics simulations showing binding of the ghrelin peptide through the extracellular TM5/TM6 entry channel of the intermediate 1 conformation through lateral diffusion. Phosphate beads are shown in green. The backbone beads of the receptor and the ghrelin peptide are depicted in pink and cyan respectively.

**S16 Figure:**
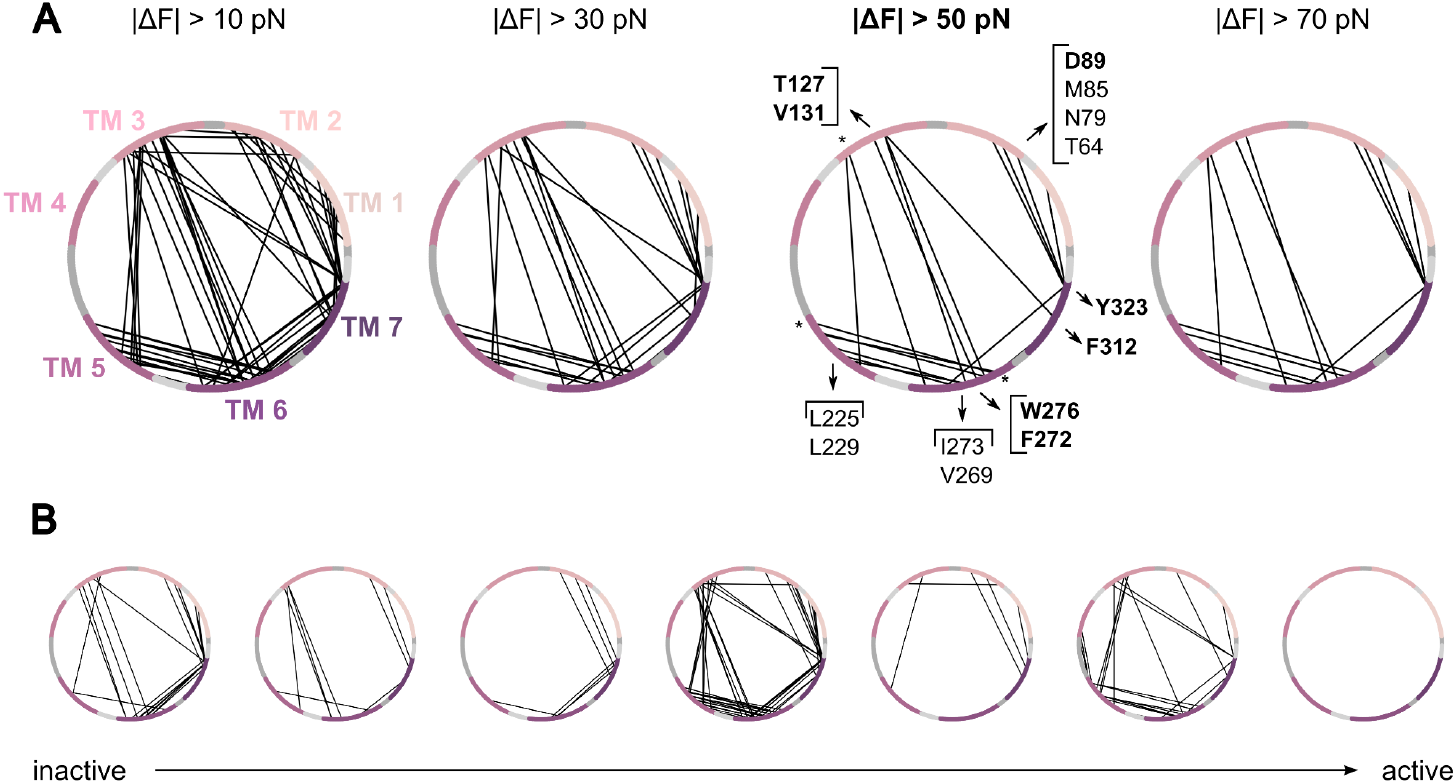
A: Change in pairwise force Δ*F*_*ij*_ = ⟨*F*_*ij*_(agonist-bound)⟩ - ⟨(*F*_*ij*_(apo)⟩ for the pair of residuse*i, j*) excluding loops at different thresholds is shown. In the 2D representations, the GHSR-1a amino acid sequence spans a circumference. Residues of the transmembrane helices are colored in a gradient starting from TM1 (pink) to TM7 (purple). The intercalated dark gray and light gray arc segments correspond to the extracellular and intracellular loop regions, respectively. Accordingly, pairs that showed a higher pairwise force in the agonist-bound state compared to the apo state are depicted as black lines. Key force transmission residues in the receptor core are labeled whereas microswitch residues are shown in bold. * denotes residue pairs that are part of the highly flexible extracellular or intracellular region of GHSR-1a. B: Differences in pairwise forces (ΔF > 50 pN) of adjacent microstates along the most probable activation path of the agonist-bound receptor. The same 2D representations were chosen as in A.

**S1 Table:**
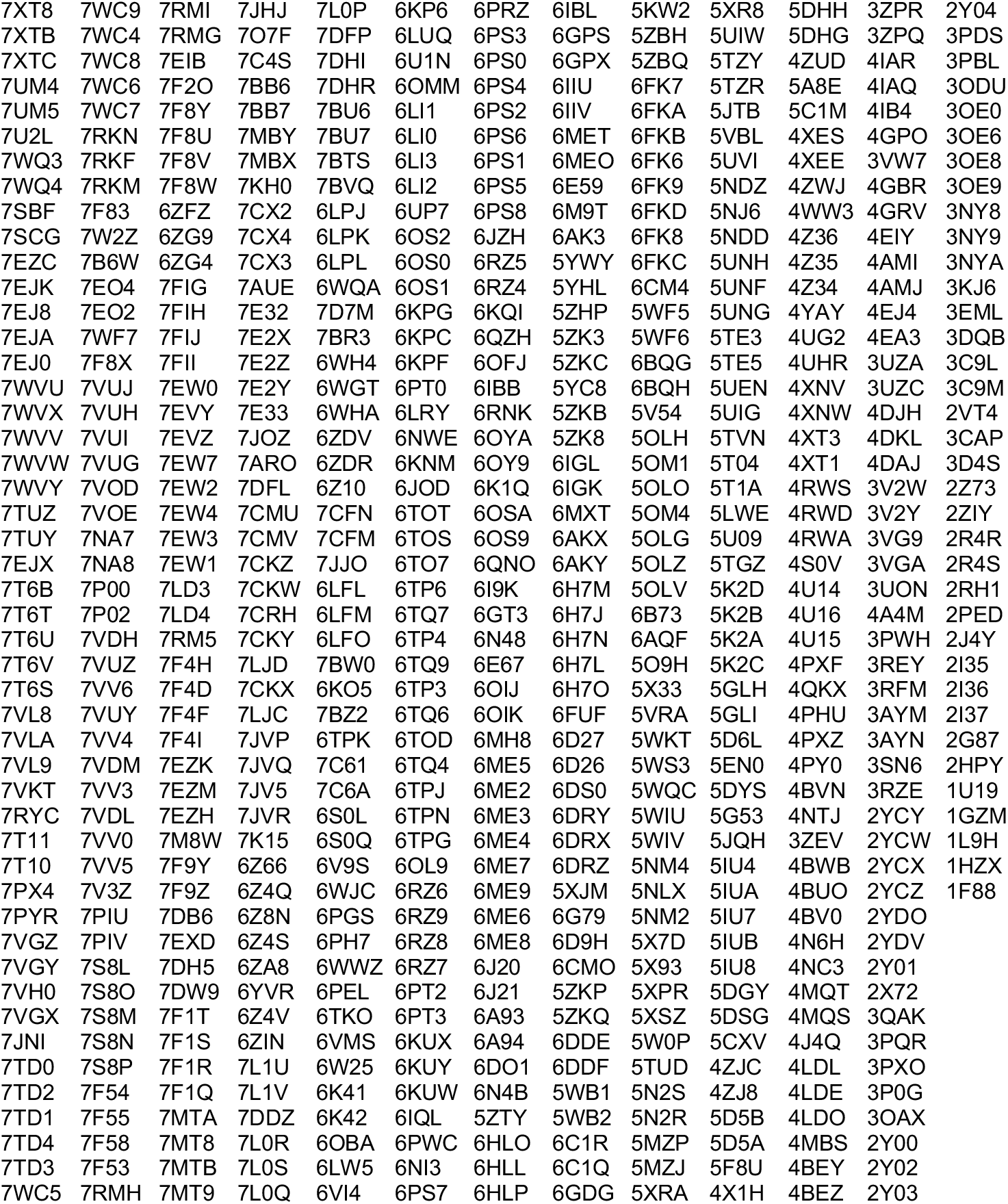
Published class A GPCR structures PDB id. (state: 11/2022) serving as input for the described computational pipeline.

