## Supplementary Information for "Unveiling G-Protein-Coupled Receptor Conformational Dynamics via Metadynamics Simulations and Markov State Models"

Rita Ann ROESSNER, Nicolas FLOQUET, Maxime LOUET*
Institut des Biomolécules Max Mousseron (IBMM), UMR5247, CNRS, Université de Montpellier, ENSCM, Pôle chimie Balard,1919 route de Mende, Montpellier, France.

### Force distribution analysis

The pair-wise force $F_{ij}$ between the pair of residues (i, j) was obtained using the force distribution analysis (FDA) tool implemented in GROMACS (version 2.11) [28]. The pair-wise force difference $\Delta F_{ij} = \langle F_{ij}(\text{agonist-bound})\rangle - \langle F_{ij}(\text{apo})\rangle$ was calculated with "agonist-bound" and "apo" indicating the respective GHSR-1a system. $\langle\rangle$ denotes an ensemble average. In practice, averages of 100 frames assigned to the lowest-energy microstate of the metastable inactive state were computed separately for both systems. The following z score function was evaluated $z = \Delta F_{ij} / \text{sqrt}[\sigma^2(F_{ij}(\text{agonist-bound})) + \sigma^2(F_{ij}(\text{apo}))]$ to give an idea of the statistical significance of the change, with σ denoting the standard deviation over the 100 frames. Pair-wise force differences that exceeded 50 pN at a z threshold of 0.75 were shown (*Fig. 4*C, *S16A* Fig.).

### Supplementary Data

**S1 Table:** Published class A GPCR structures PDB id. (state: 11/2022) serving as input for the described computational pipeline.

| | | | | | | | | | | | | |
|---|---|---|---|---|---|---|---|---|---|---|---|---|
| 7XT8 | 7WC9 | 7RMI | 7JHJ | 7L0P | 6KP6 | 6PRZ | 6IBL | 5KW2 | 5XR8 | 5DHH | 3ZPR | 2Y04 |
| 7XTB | 7WC4 | 7RMG | 7O7F | 7DFP | 6LUQ | 6PS3 | 6GPS | 5ZBH | 5UIW | 5DHG | 3ZPQ | 3PDS |
| 7XTC | 7WC8 | 7EIB | 7C4S | 7DHI | 6U1N | 6PS0 | 6GPX | 5ZBQ | 5TZY | 4ZUD | 4IAR | 3PBL |
| 7UM4 | 7WC6 | 7F2O | 7BB6 | 7DHR | 6OMM | 6PS4 | 6IIU | 6FK7 | 5TZR | 5A8E | 4IAQ | 3ODU |
| 7UM5 | 7WC7 | 7F8Y | 7BB7 | 7BU6 | 6LI1 | 6PS2 | 6IIV | 6FKA | 5JTB | 5C1M | 4IB4 | 3OE0 |
| 7U2L | 7RKN | 7F8U | 7MBY | 7BU7 | 6LI0 | 6PS6 | 6MET | 6FKB | 5VBL | 4XES | 4GPO | 3OE6 |
| 7WQ3 | 7RKF | 7F8V | 7MBX | 7BTS | 6LI3 | 6PS1 | 6MEO | 6FK6 | 5UVI | 4XEE | 3VW7 | 3OE8 |
| 7WQ4 | 7RKM | 7F8W | 7KH0 | 7BVQ | 6LI2 | 6PS5 | 6E59 | 6FK9 | 5NDZ | 4ZWJ | 4GBR | 3OE9 |
| 7SBF | 7F83 | 6ZFZ | 7CX2 | 6LPJ | 6UP7 | 6PS8 | 6M9T | 6FKD | 5NJ6 | 4WW3 | 4GRV | 3NY8 |
| 7SCG | 7W2Z | 6ZG9 | 7CX4 | 6LPK | 6OS2 | 6JZH | 6AK3 | 6FK8 | 5NDD | 4Z36 | 4EIY | 3NY9 |
| 7EZC | 7B6W | 6ZG4 | 7CX3 | 6LPL | 6OS0 | 6RZ5 | 5YWY | 6FKC | 5UNH | 4Z35 | 4AMI | 3NYA |
| 7EJK | 7EO4 | 7FIG | 7AUE | 6WQA | 6OS1 | 6RZ4 | 5YHL | 6CM4 | 5UNF | 4Z34 | 4AMJ | 3KJ6 |
| 7EJ8 | 7EO2 | 7FIH | 7E32 | 7D7M | 6KPG | 6KQI | 5ZHP | 5WF5 | 5UNG | 4YAY | 4EJ4 | 3EML |
| 7EJA | 7WF7 | 7FIJ | 7E2X | 7BR3 | 6KPC | 6QZH | 5ZK3 | 5WF6 | 5TE3 | 4UG2 | 4EA3 | 3DQB |
| 7EJ0 | 7F8X | 7FII | 7E2Z | 6WH4 | 6KPF | 6OFJ | 5ZKC | 6BQG | 5TE5 | 4UHR | 3UZA | 3C9L |
| 7WVU | 7VUJ | 7EW0 | 7E2Y | 6WGT | 6PT0 | 6IBB | 5YC8 | 6BQH | 5UEN | 4XNV | 3UZC | 3C9M |
| 7WVX | 7VUH | 7EVY | 7E33 | 6WHA | 6LRY | 6RNK | 5ZKB | 5V54 | 5UIG | 4XNW | 4DJH | 2VT4 |
| 7WVV | 7VUI | 7EVZ | 7JOZ | 6ZDV | 6NWE | 6OYA | 5ZK8 | 5OLH | 5TVN | 4XT3 | 4DKL | 3CAP |
| 7WVW | 7VUG | 7EW7 | 7ARO | 6ZDR | 6KNM | 6OY9 | 6IGL | 5OM1 | 5T04 | 4XT1 | 4DAJ | 3D4S |
| 7WVY | 7VOD | 7EW2 | 7DFL | 6Z10 | 6JOD | 6K1Q | 6IGK | 5OLO | 5T1A | 4RWS | 3V2W | 2Z73 |
| 7TUZ | 7VOE | 7EW4 | 7CMU | 7CFN | 6TOT | 6OSA | 6MXT | 5OM4 | 5LWE | 4RWD | 3V2Y | 2ZIY |
| 7TUY | 7NA7 | 7EW3 | 7CMV | 7CFM | 6TOS | 6OS9 | 6AKX | 5OLG | 5U09 | 4RWA | 3VG9 | 2R4R |
| 7EJX | 7NA8 | 7EW1 | 7CKZ | 7JJO | 6TO7 | 6QNO | 6AKY | 5OLZ | 5TGZ | 4S0V | 3VGA | 2R4S |
| 7T6B | 7P00 | 7LD3 | 7CKW | 6LFL | 6TP6 | 6I9K | 6H7M | 5OLV | 5K2D | 4U14 | 3UON | 2RH1 |
| 7T6T | 7P02 | 7LD4 | 7CRH | 6LFM | 6TQ7 | 6GT3 | 6H7J | 6B73 | 5K2B | 4U16 | 4A4M | 2PED |
| 7T6U | 7VDH | 7RM5 | 7CKY | 6LFO | 6TP4 | 6N48 | 6H7N | 6AQF | 5K2A | 4U15 | 3PWH | 2J4Y |
| 7T6V | 7VUZ | 7F4H | 7LJD | 7BW0 | 6TQ9 | 6E67 | 6H7L | 5O9H | 5K2C | 4PXF | 3REY | 2I35 |
| 7T6S | 7VV6 | 7F4D | 7CKX | 6KO5 | 6TP3 | 6OIJ | 6H7O | 5X33 | 5GLH | 4QKX | 3RFM | 2I36 |
| 7VL8 | 7VUY | 7F4F | 7LJC | 7BZ2 | 6TQ6 | 6OIK | 6FUF | 5VRA | 5GLI | 4PHU | 3AYM | 2I37 |
| 7VLA | 7VV4 | 7F4I | 7JVP | 6TPK | 6TOD | 6MH8 | 6D27 | 5WKT | 5D6L | 4PXZ | 3AYN | 2G87 |
| 7VL9 | 7VDM | 7EZK | 7JVQ | 7C61 | 6TQ4 | 6ME5 | 6D26 | 5WS3 | 5EN0 | 4PY0 | 3SN6 | 2HPY |
| 7VKT | 7VV3 | 7EZM | 7JV5 | 7C6A | 6TPJ | 6ME2 | 6DS0 | 5WQC | 5DYS | 4BVN | 3RZE | 1U19 |
| 7RYC | 7VDL | 7EZH | 7JVR | 6S0L | 6TPN | 6ME3 | 6DRY | 5WIU | 5G53 | 4NTJ | 2YCY | 1GZM |
| 7T11 | 7VV0 | 7M8W | 7K15 | 6S0Q | 6TPG | 6ME4 | 6DRX | 5WIV | 5JQH | 3ZEV | 2YCW | 1L9H |
| 7T10 | 7VV5 | 7F9Y | 6Z66 | 6V9S | 6OL9 | 6ME7 | 6DRZ | 5NM4 | 5IU4 | 4BWB | 2YCX | 1HZX |
| 7PX4 | 7V3Z | 7F9Z | 6Z4Q | 6WJC | 6RZ6 | 6ME9 | 5XJM | 5NLX | 5IUA | 4BUO | 2YCZ | 1F88 |
| 7PYR | 7PIU | 7DB6 | 6Z8N | 6PGS | 6RZ9 | 6ME6 | 6G79 | 5NM2 | 5IU7 | 4BV0 | 2YDO | |
| 7VGZ | 7PIV | 7EXD | 6Z4S | 6PH7 | 6RZ8 | 6ME8 | 6D9H | 5X7D | 5IUB | 4N6H | 2YDV | |
| 7VGY | 7S8L | 7DH5 | 6ZA8 | 6WWZ | 6RZ7 | 6J20 | 6CMO | 5X93 | 5IU8 | 4NC3 | 2Y01 | |
| 7VH0 | 7S8O | 7DW9 | 6YVR | 6PEL | 6PT2 | 6J21 | 5ZKP | 5XPR | 5DGY | 4MQT | 2X72 | |
| 7VGX | 7S8M | 7F1T | 6Z4V | 6TKO | 6PT3 | 6A93 | 5ZKQ | 5XSZ | 5DSG | 4MQS | 3QAK | |
| 7JNI | 7S8N | 7F1S | 6ZIN | 6VMS | 6KUX | 6A94 | 6DDE | 5W0P | 5CXV | 4J4Q | 3PQR | |
| 7TD0 | 7S8P | 7F1R | 7L1U | 6W25 | 6KUY | 6DO1 | 6DDF | 5TUD | 4ZJC | 4LDL | 3PXO | |
| 7TD2 | 7F54 | 7F1Q | 7L1V | 6K41 | 6KUW | 6N4B | 5WB1 | 5N2S | 4ZJ8 | 4LDE | 3P0G | |
| 7TD1 | 7F55 | 7MTA | 7DDZ | 6K42 | 6IQL | 5ZTY | 5WB2 | 5N2R | 5D5B | 4LDO | 3OAX | |
| 7TD4 | 7F58 | 7MT8 | 7L0R | 6OBA | 6PWC | 6HLO | 6C1R | 5MZP | 5D5A | 4MBS | 2Y00 | |
| 7TD3 | 7F53 | 7MTB | 7L0S | 6LW5 | 6NI3 | 6HLL | 6C1Q | 5MZJ | 5F8U | 4BEY | 2Y02 | |
| 7WC5 | 7RMH | 7MT9 | 7L0Q | 6VI4 | 6PS7 | 6HLP | 6GDG | 5XRA | 4X1H | 4BEZ | 2Y03 | |

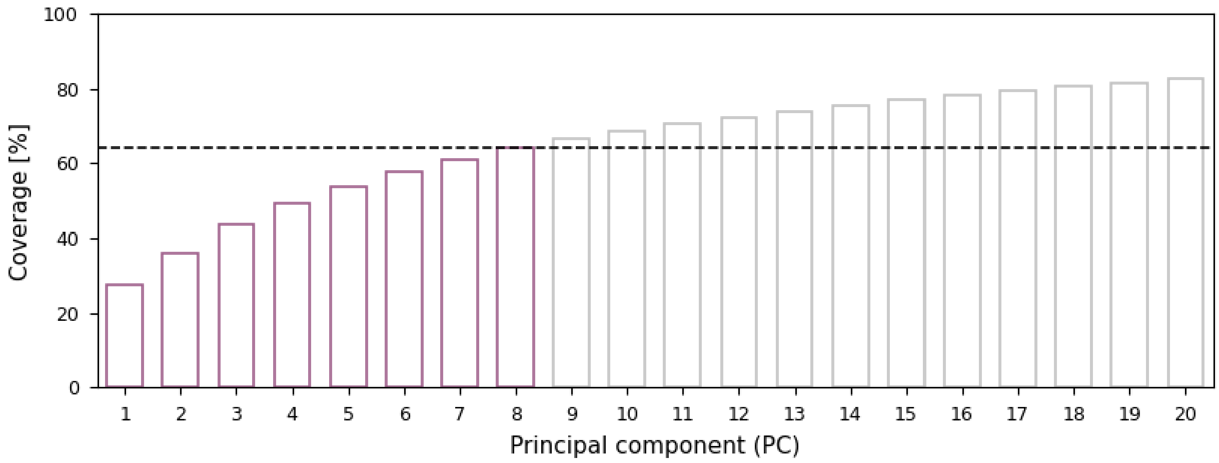


**S2 Figure:** Percentage of the cumulative variance described by PCA as a function of the respective PCs. The dashed line indicates the value for the eight PCs (64 %) that were considered as collective variables for the metadynamics simulations.

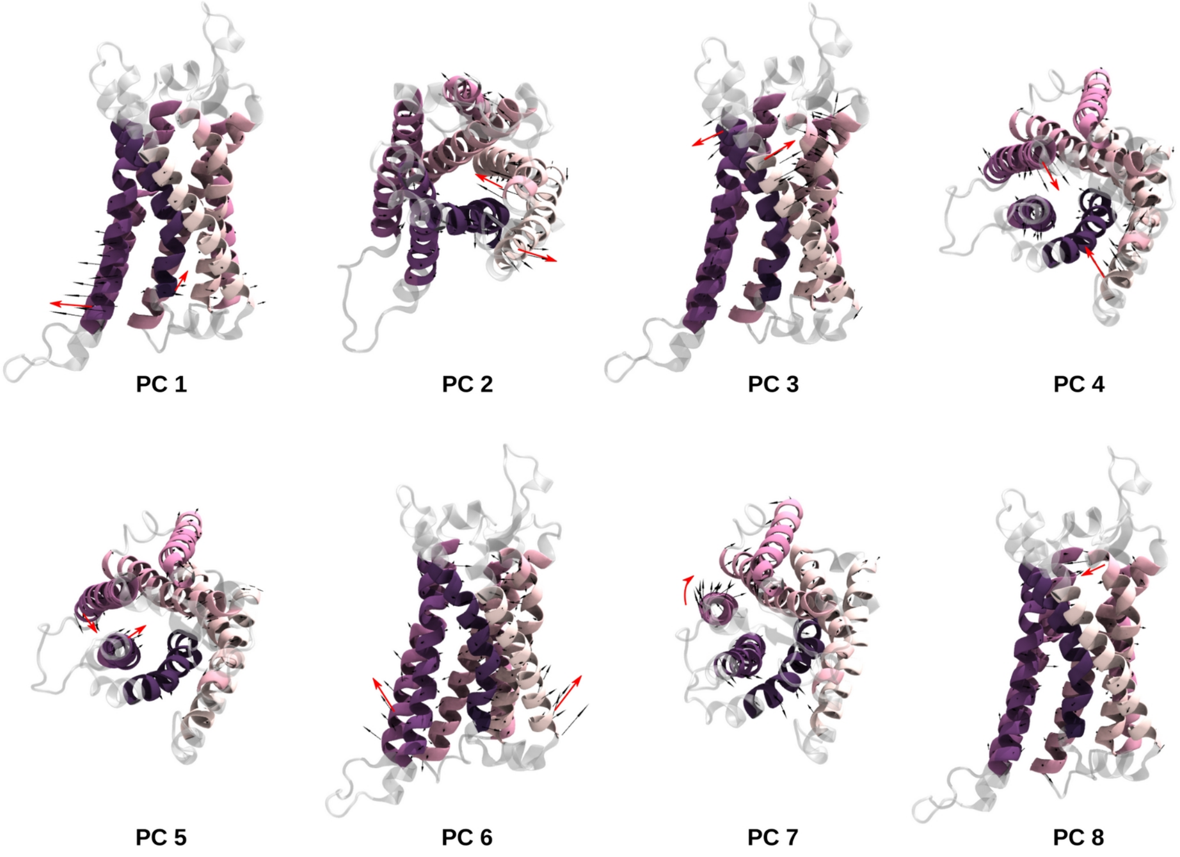


**S3 Figure:** Collective motions along the first eight principal components. GHSR-1a is shown in cartoon representation. The residues of the membrane-spanning region considered in PCA are colored by index (pink to purple); the remaining residues are shown in gray. The black arrows indicate the collective motion along the respective PC, with the arrow length corresponding to the amplitude of the vector. The red arrows highlight the largest rearrangements.

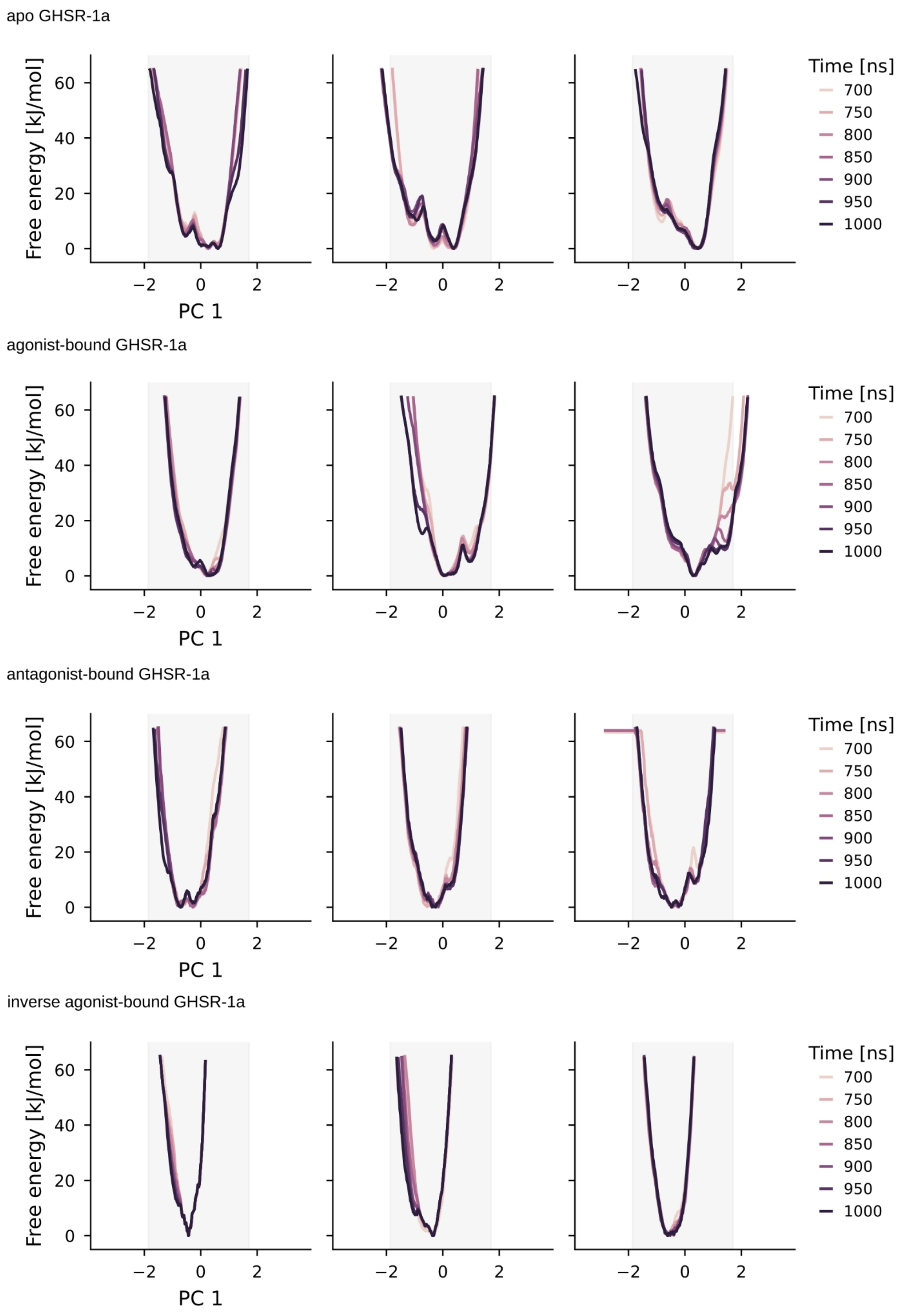


**S4 Figure:** Time dependence of calculated free energy profiles obtained from metadynamics simulations. Traces as a function of PC1 are visualized for all n=3 replicas of apo, agonist-bound, antagonist ghrelin-bound, and inverse agonist-bound systems. The color gradient from light to dark indicates the progress of the simulations. The profiles remain unchanged over the last 300 ns, indicating that the simulations have converged.

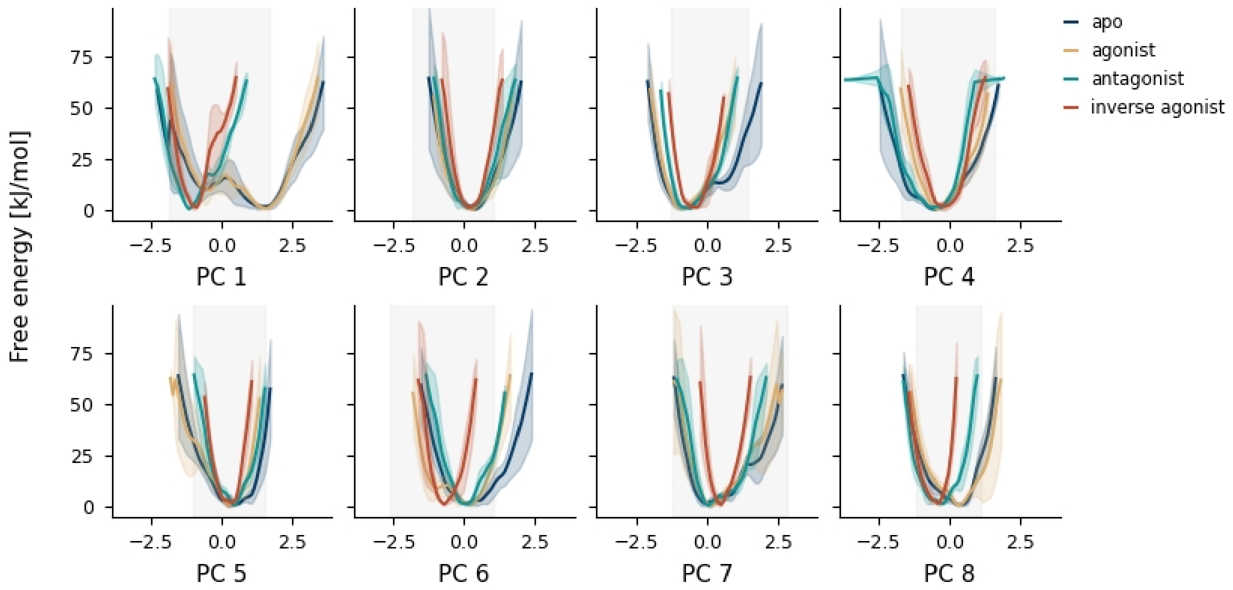


**S5 Figure:** Free energy profiles obtained from metadynamics simulations. Average ± standard deviation of free energy profiles (n=3 independent replicas) as a function of PC1 to PC8 for the apo- (green), ghrelin-bound, (yellow), antagonist-bound (green), and inverse agonist system (red). The area in which the projections of the experimental structures are found is depicted in light grey.

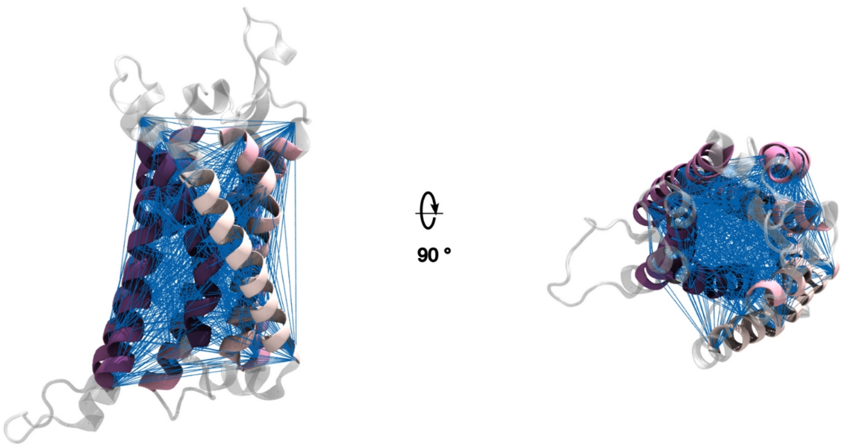


**S6 Figure:** Cα distance features used as TICA input. GHSR-1a is shown in cartoon representation. The residues of the membrane-spanning region considered in PCA are colored by index (pink to purple); the remaining residues are shown in gray. The distances between the Cα atoms that we used for TICA and subsequent MSM construction are depicted as blue lines.

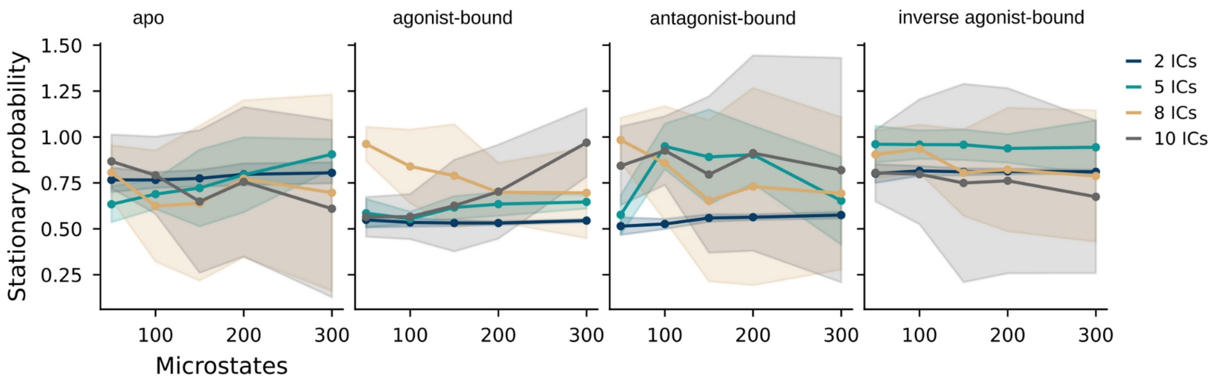


**S7 Figure:** Optimization of MSM hyperparameters. The VAMP2 scores are calculated at varying numbers of microstates and TICA dimensions (ICs) for the apo, agonist-bound, antagonist-bound, and inverse agonist-bound systems. The mean values from five-fold cross-validation are plotted as dots and standard deviations as shaded regions.

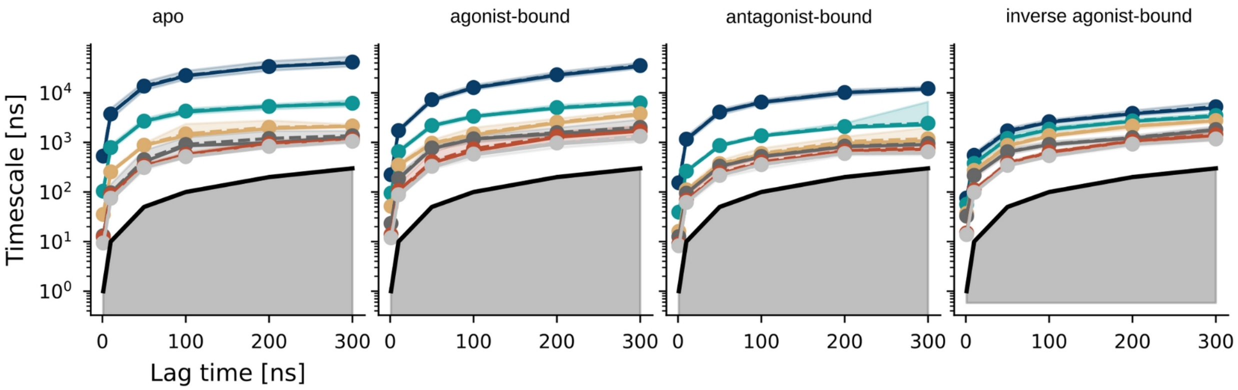


**S8 Figure:** Implied timescale analysis to identify a memoryless Markovian time. The top six eigenvalues of the transition probability matrix for the apo, agonist-bound, antagonist-bound, and inverse agonist-bound systems are calculated at varying lag times. The solid lines correspond to the implied time scales of maximum likelihood MSMs. The 95% confidence intervals, quantified based on a Bayesian scheme, are depicted as shaded regions. The sample means are given by dashed lines. The black solid curve indicates the time resolution limit of the estimated Markov model.

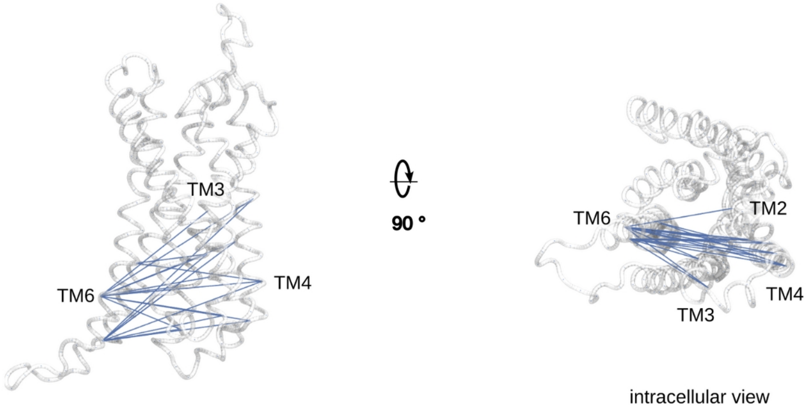


**S9 Figure:** Side and intracellular view of GHSR-1a represented as ribbons with the 20 Cα-distances exhibiting the largest correlation with IC 1 highlighted in blue.

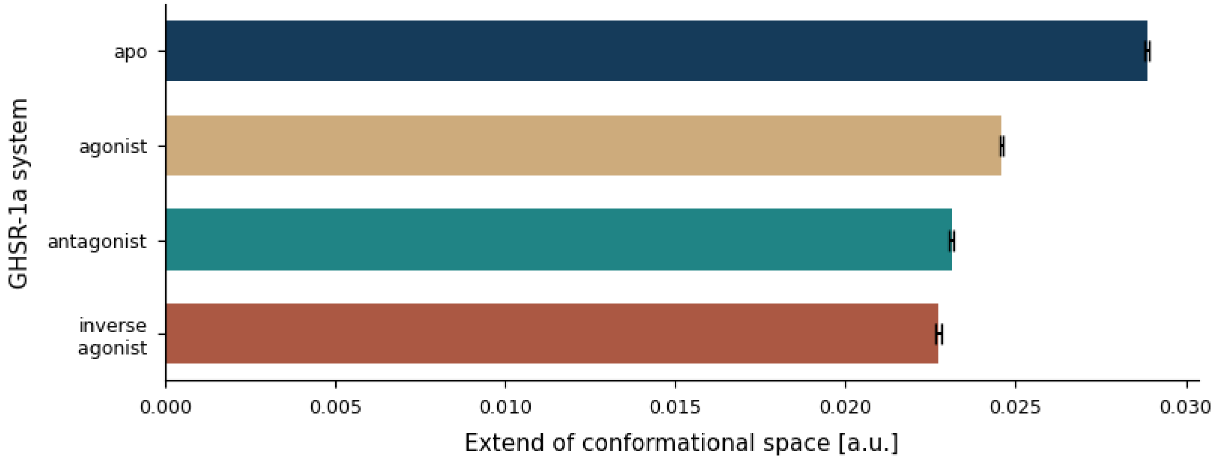


**S10 Figure:** Extend of the conformational space explored by each system during unbiased MD simulations on the basis of which the MSMs were constructed. We calculated the Pearson correlation coefficient of the Cα-distance features between each trajectory frame used for MSM construction and a reference (PDB id. 7F9Y). We calculated the extent of the correlation value distribution, excluding outliers whereby the values presented in the barplot indicate the mean $\pm$ standard deviation obtained from bootstrapping using 1000 samples.

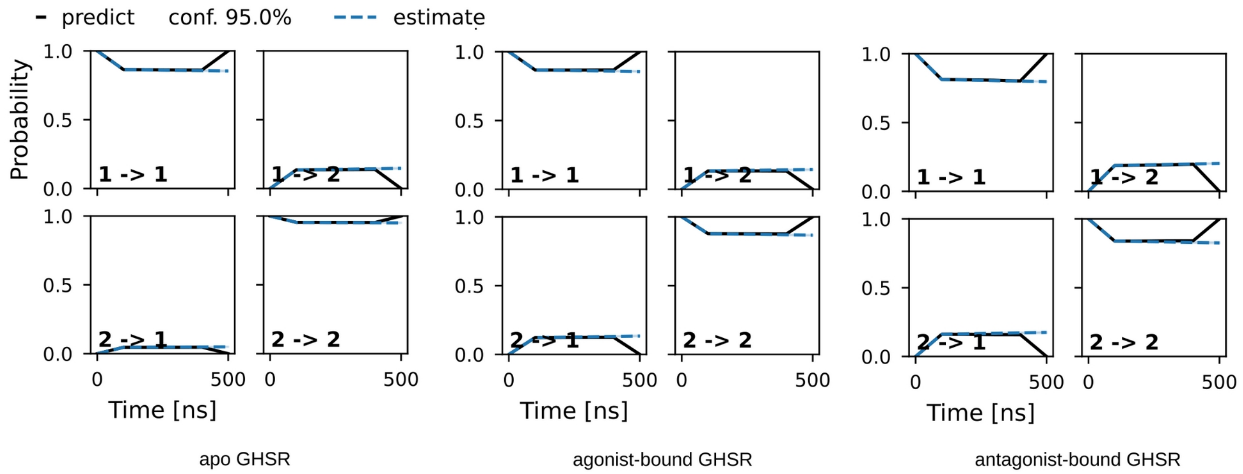


**S11 Figure:** Chapman-Kolmogorov test to assess the kinetic self-consistency of the MSM. The tests for the apo, agonist-bound, and antagonist-bound systems indicate that the model predictions (black line) at long time scales are consistent with the corresponding estimates (blue line). Estimates are shown with 95% confidence intervals.

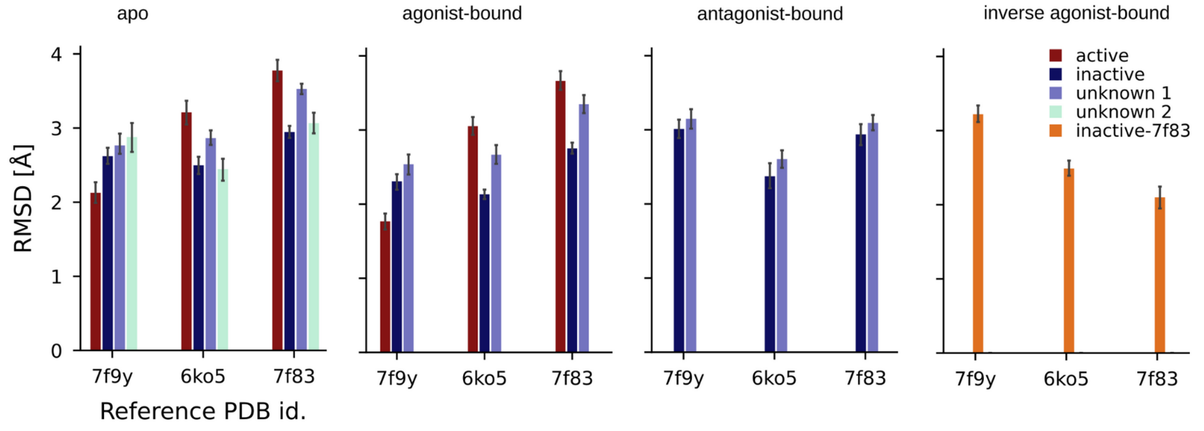


**S12 Figure:** RMSD (average ± standard deviation) of the lowest energy models of each macrostate of all GHSR-1a systems with respect to the experimental structures of the agonist- (PDB id. 7F9Y) antagonist- (PDB id. 6KO5) and inverse agonist-bound receptor (PDB id. 7F83).

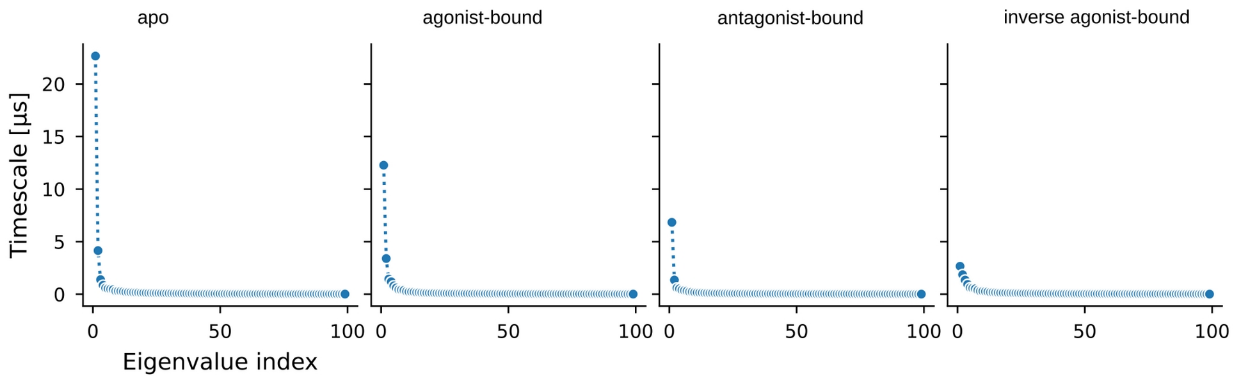


**S13 Figure:** Spectral analysis of the eigenvalues of the apo, agonist-bound, antagonist-bound, and inverse agonist-bound system to identify the number of PCCA+ clusters.

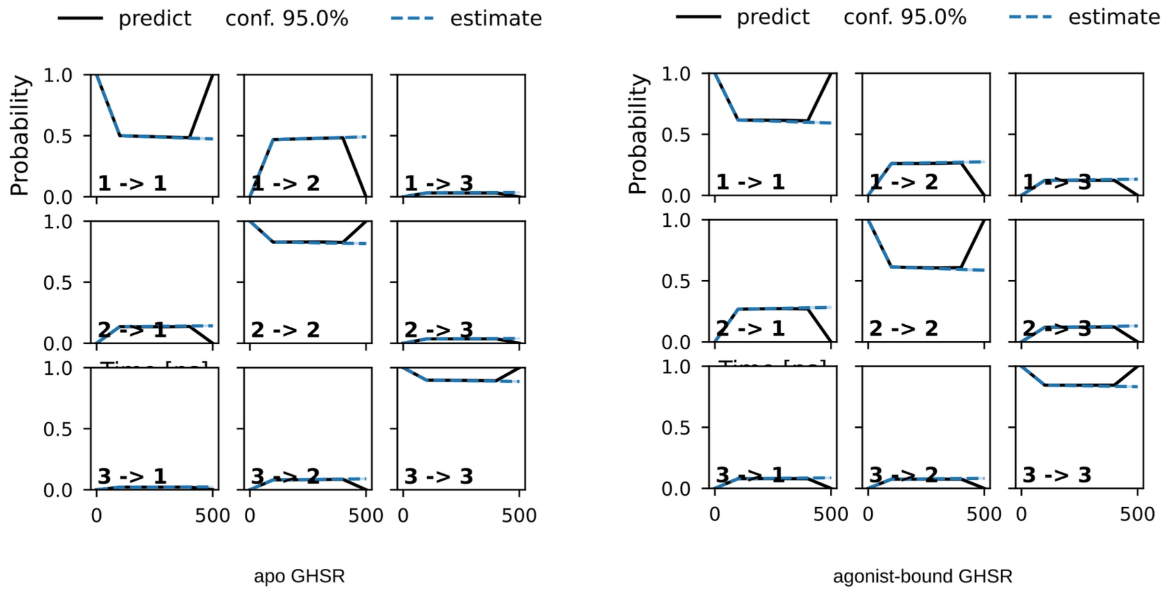


**S14 Figure:** Chapman-Kolmogorov test to assess the kinetic self-consistency of the MSM. The tests for the apo, and agonist-bound systems indicate that the model predictions (black line) at long time scales are consistent with the corresponding estimates (blue line). Estimates are shown with 95% confidence intervals.

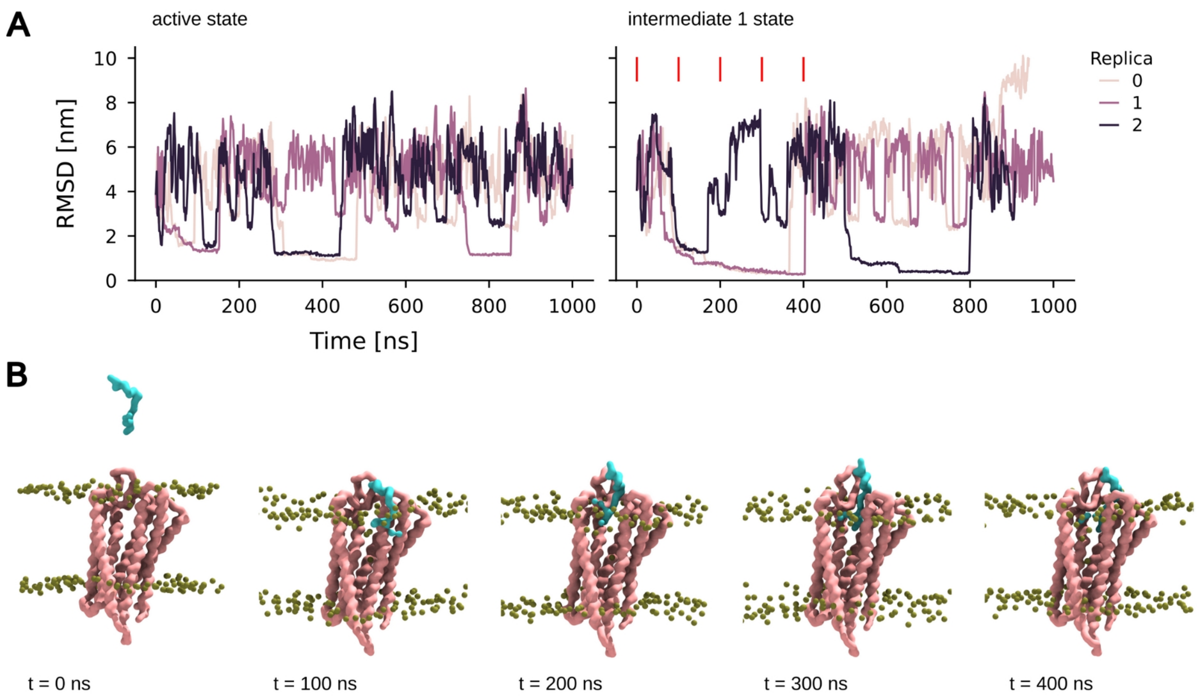


**S15 Figure:** Ghrelin binding to GHSR-1a in intermediate 1 state in CG metadynamics simulations. A: Time traces of the RMSD computed ghrelin peptide backbone bead compared to the coarse-grained reference structure of the ghrelin-bound receptor (PDB id. 7F9Y) along the metadynamics simulations of GHSR-1a in its active (left) or intermediate 1 conformer (right). Different colors correspond to different replicas (n=3 per system). The red lines indicate the time points of the snapshots in B taken from replica 1 of the intermediate 1 simulations. B: Snapshot along the CG metadynamics simulations showing binding of the ghrelin peptide through the extracellular TM5/TM6 entry channel of the intermediate 1 conformation through lateral diffusion. Phosphate beads are shown in green. The backbone beads of the receptor and the ghrelin peptide are depicted in pink and cyan respectively.

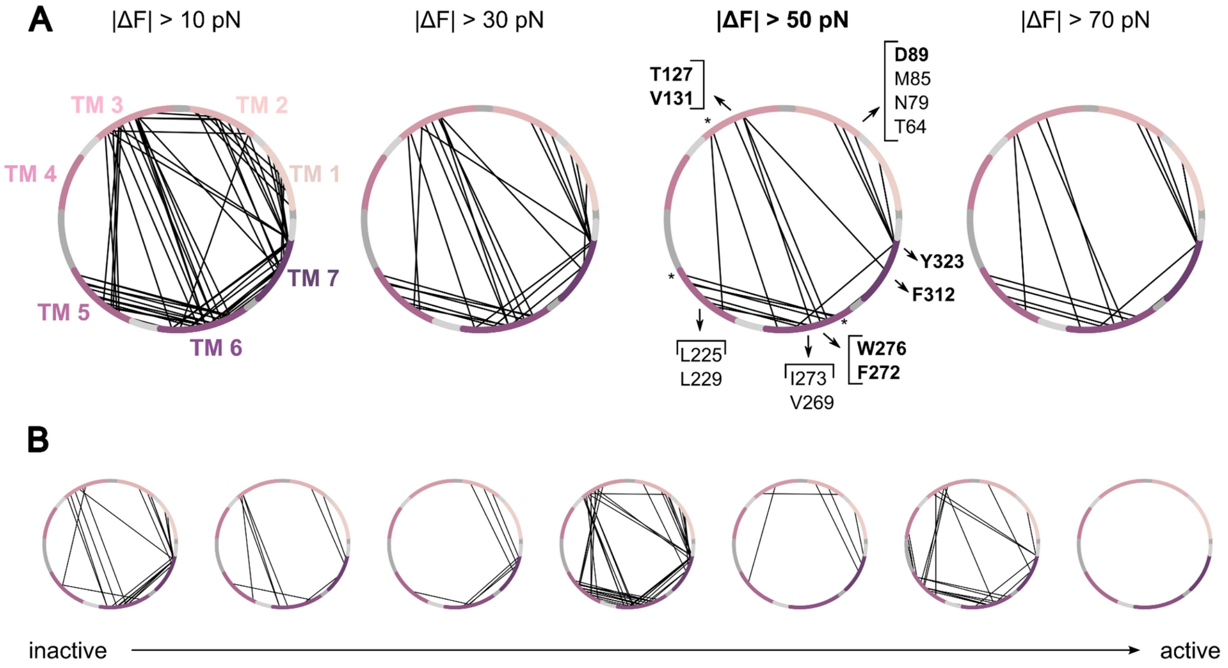


**S16 Figure:** A: Change in pairwise force $\Delta F_{ij} = \langle F_{ij}(\text{agonist-bound})\rangle - \langle F_{ij}(\text{apo})\rangle$ for the pair of residues ($i$, $j$) excluding loops at different thresholds is shown. In the 2D representations, the GHSR-1a amino acid sequence spans a circumference. Residues of the transmembrane helices are colored in a gradient starting from TM1 (pink) to TM7 (purple). The intercalated dark gray and light gray arc segments correspond to the extracellular and intracellular loop regions, respectively. Accordingly, pairs that showed a higher pairwise force in the agonist-bound state compared to the apo state are depicted as black lines. Key force transmission residues in the receptor core are labeled whereas microswitch residues are shown in bold. * denotes residue pairs that are part of the highly flexible extracellular or intracellular region of GHSR-1a. B: Differences in pairwise forces (ΔF > 50 pN) of adjacent microstates along the most probable activation path of the agonist-bound receptor. The same 2D representations were chosen as in A.
